# Large-scale phylogenomics of the genus *Macrostomum* (Platyhelminthes) reveals cryptic diversity and novel sexual traits

**DOI:** 10.1101/2021.03.28.437366

**Authors:** Jeremias N. Brand, Gudrun Viktorin, R. Axel W. Wiberg, Christian Beisel, Lukas Schärer

## Abstract

Free-living flatworms of the genus *Macrostomum* are small and transparent animals, representing attractive study organisms for a broad range of topics in evolutionary, developmental, and molecular biology. The genus includes the model organism *M. lignano* for which extensive molecular resources are available, and recently there is a growing interest in extending work to additional species in the genus. These endeavours are currently hindered because, even though >200 *Macrostomum* species have been taxonomically described, molecular phylogenetic information and geographic sampling remain limited. We report on a global sampling campaign aimed at increasing taxon sampling and geographic representation of the genus. Specifically, we use extensive transcriptome and single-locus data to generate phylogenomic hypotheses including 145 species. Across different phylogenetic methods and alignments used, we identify several consistent clades, while their exact grouping is less clear, possibly due to a radiation early in *Macrostomum* evolution. Moreover, we uncover a large undescribed diversity, with 94 of the studied species likely being new to science, and we identify multiple novel morphological traits. Furthermore, we identify cryptic speciation in a taxonomically challenging assemblage of species, suggesting that the use of molecular markers is a prerequisite for future work, and we describe the distribution of possible synapomorphies and suggest taxonomic revisions based on our finding. Our large-scale phylogenomic dataset now provides a robust foundation for comparative analyses of morphological, behavioural and molecular evolution in this genus.

## 1. Introduction

The genus *Macrostomum* (Macrostomorpha, Platyhelminthes) is a large clade of free-living flatworms with a global distribution in marine, brackish and freshwater environments (Ferguson 1954). After the Catenulida, the Macrostomorpha are the earliest-branching lineage of Platyhelminthes (Laumer et al. 2015), giving *Macrostomum* an interesting taxonomic position, and representing a valuable point of comparison to other well-studied flatworms such as members of the Tricladida and the parasitic Neodermata. The genus contains the model organism *M. lignano* (Ladurner et al. 2005; Wudarski et al. 2020), which, in part due to its high transparency and experimental amenability, is used for the study of a variety of biological topics, including aging (Mouton et al. 2018), bioadhesion (Lengerer et al. 2016; Wunderer et al. 2019), regeneration and stem cells (Egger et al. 2006; Grudniewska et al. 2016; Lengerer et al. 2016), and sexual selection (Schärer et al. 2011; Marie-Orleach et al. 2020). A rich experimental and genomic toolset is available for *M. lignano*, including *in situ* hybridization, RNA interference, transgenesis, a range of transcriptomes, and a sequenced genome (Pfister et al. 2007, 2008; Wasik et al. 2015; Wudarski et al. 2017). Perhaps as a result of the emergence of *Macrostomum* as a promising model organism, there has been a resurgence of taxonomic research on this genus and related taxa (e.g. Ladurner et al. 2005; Adami et al. 2012; Janssen et al. 2015; Sun et al. 2015; Atherton and Jondelius 2019; Xin et al. 2019; Schärer et al. 2020), after an initially very active period of research in the middle of the last century (e.g. Luther 1947, 1960; Ferguson 1954).

Moreover, primarily due to the discovery of a whole-genome duplication and accompanying karyotype instability in *M. lignano* (Zadesenets et al. 2016, 2017), there is increased interest in expanding the genomic resources and molecular phylogenetic placement of additional species in the genus,. To this end, recent work has provided transcriptomic resources for additional species (Brand et al. 2020) and field-collections have revealed similar karyological rearrangements to the ones observed in *M. lignano* in some closely related species, while others were found to have stable diploid genomes (Schärer et al. 2020; Zadesenets et al. 2020). Accurate phylogenetic information will greatly help guide the search for additional *Macrostomum* model species and permit to study the (co)evolution of the large diversity of morphological and behavioural traits in the genus, a brief introduction of which we provide in the following.

Members of the genus *Macrostomum* have intriguing aflagellate sperm that in many species carry two stiff lateral bristles and a terminal brush (Fig. 1A), while other species have sperm that lack these structures (Fig. 1B). Moreover, species show a large diversity in both the morphology of the stylet (Fig. 2) and the female antrum. Previous work has also shown that there are two main clades of *Macrostomum* that differ in terms of their mating behaviour (Schärer et al. 2011). One clade (called “Clade I” in Schärer et al. 2011) was characterized by a stereotypical stylet and sperm morphology, and species in that clade are thought to mate exclusively through hypodermic insemination, a form of traumatic mating where the ejaculate is injected though the mating partner’s epidermis into the tissue using a needle-like copulatory organ (Fig. 1B). The second clade (called “Clade II” in Schärer et al. 2011) contains species that are thought to mate primarily through reciprocal copulation, where both individuals reciprocally insert their (often blunt) stylet (Fig. 1A) into the female antrum of the mating partner, and are able to donate and receive ejaculate at the same time (see also Schärer et al. 2004, 2011; Singh et al. 2020). Within Clade II one species, *M. hystrix*, was found to have convergently evolved the hypodermic mating behaviour, and this was associated with convergent changes in stylet, sperm, and antrum morphology, leading to a morphology that is very similar to that of *M. pusillum* (Fig. 1B).

**Fig. 1.**
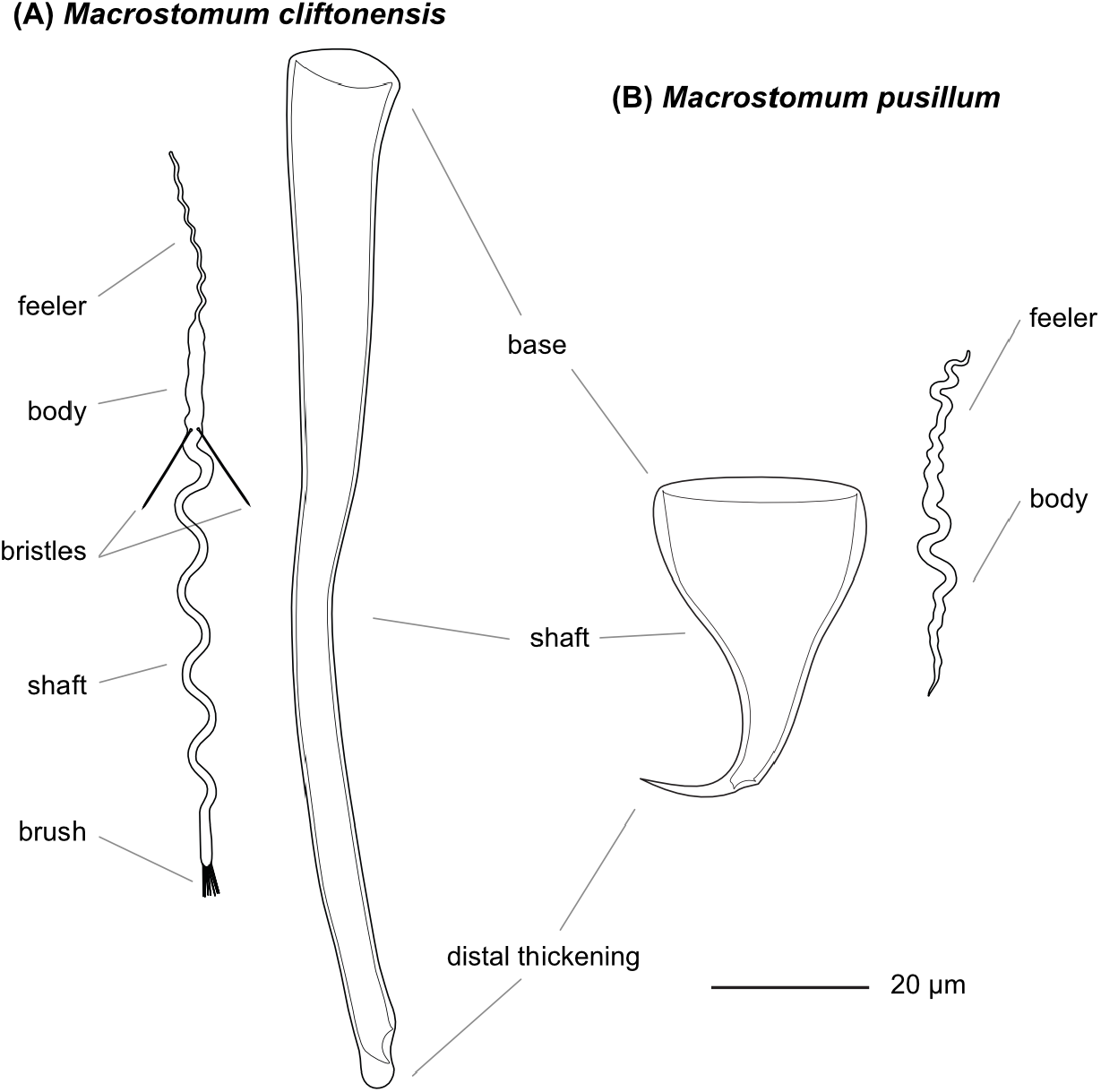
Sperm and stylet morphology of two *Macrostomum* species. (A) *M. cliftonensis*, a species that represents the reciprocal mating syndrome, with sperm carrying bristles and a brush, and a stylet with a blunt distal thickening and (B) *M. pusillum*, a species that represents the hypodermic mating syndrome, with simple sperm and a needle-like stylet.

**Fig. 2.**
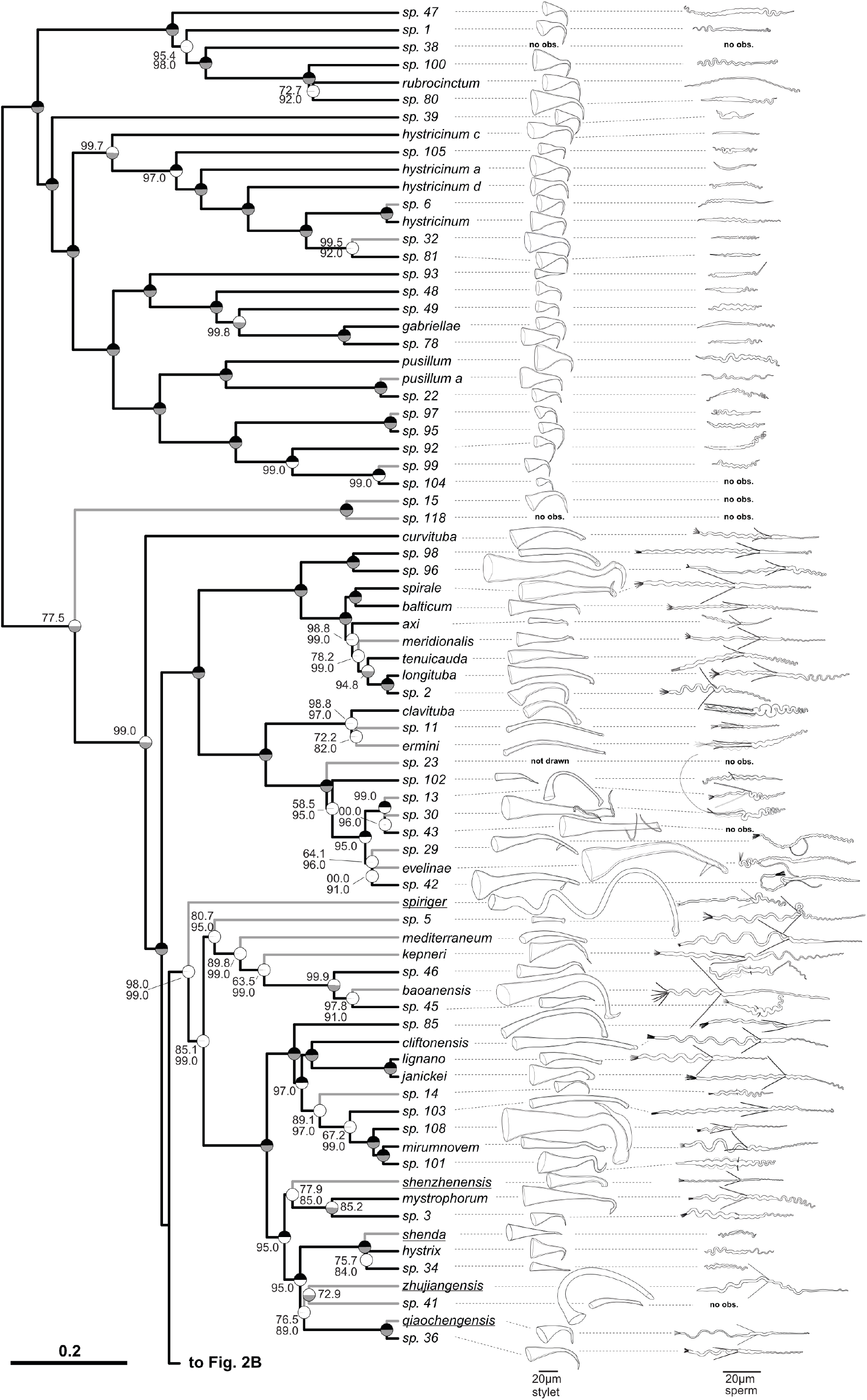
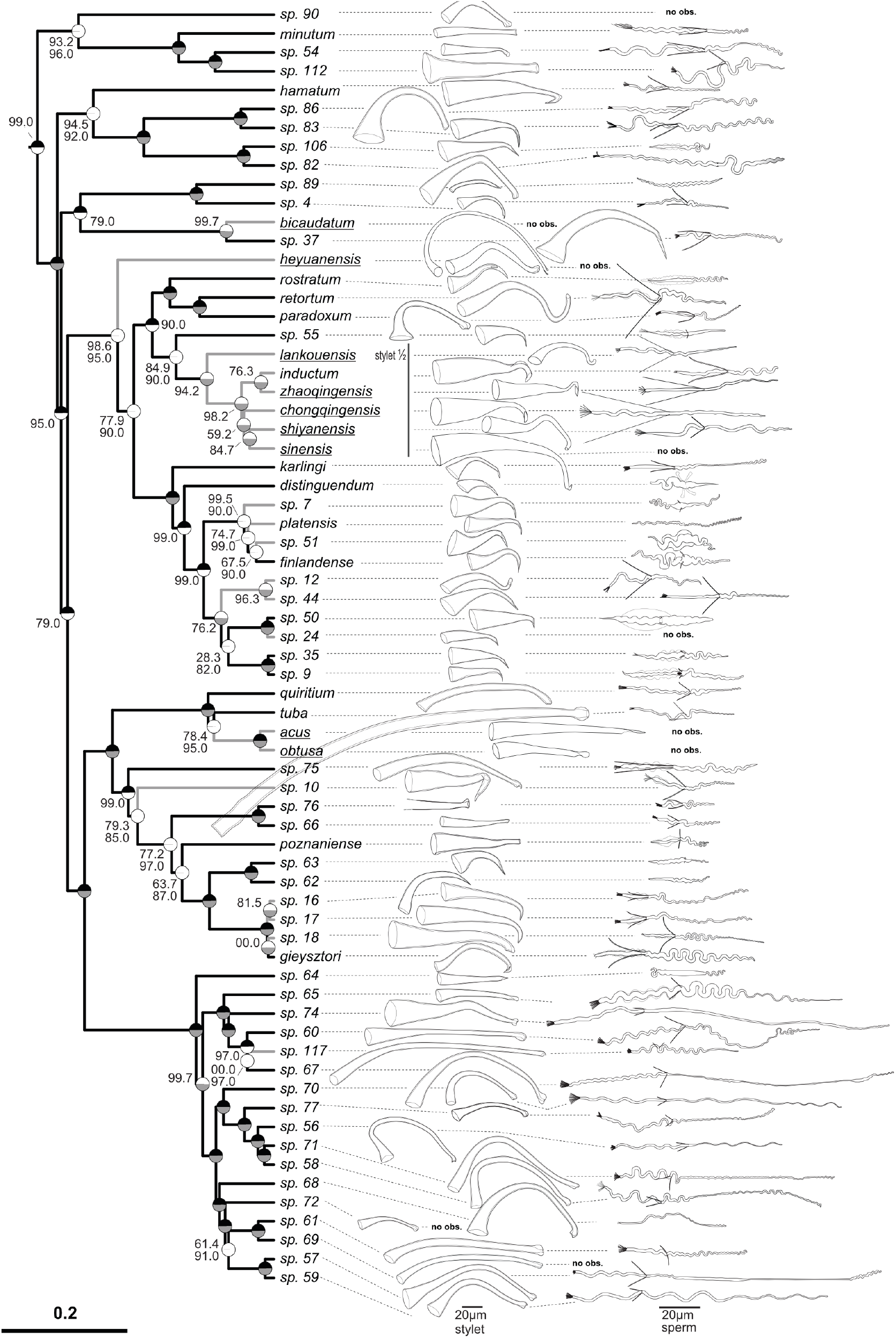
Phylogeny of the genus *Macrostomum*, showing the striking diversity in stylet and sperm morphology across the genus. The ultrametric phylogeny (C-IQ-TREE) includes all 145 studied species (with 77 species depicted in Fig. 2A and 68 species in Fig. 2B), while additional phylogenies based solely on transcriptomes are shown in Fig. 3 and Fig. S2. Branch supports are ultrafast bootstraps (top, black if 100) and approximate likelihood ratio tests (bottom, grey if 100). Species without available transcriptomes that were included in C-IQ-TREE based on a *28S rRNA* fragment are indicated with grey branches. Stylet and sperm are drawn based on our live observations, except for species with underlined names, which were redrawn based on the species descriptions (*M. acus, M. obtusa* and *M. sinensis* from Wang 2005; *M. heyuanensis* and *M. bicaudatum* from Sun et al. 2015; *M. chongqingensis* and *M. zhaoqingensis* from Lin et al. 2017a; *M. shiyanensis* and *M. lankouensis* from Lin et al. 2017b; *M. shenzhenensis* and *M. qiaochengensis* from Wang et al. 2017; and *M. spiriger* and *M. shenda* from Xin et al. 2019). Note that the stylet of *M*. sp. 15 is not drawn to scale, the stylets of some species are drawn at half size (stylet ½), and the stylet of *M*. sp. 23 is not drawn since it was incomplete (specimen ID MTP LS 913). Unobserved structures are marked as no observation (no obs.).

Understanding the molecular phylogeny of the genus will furthermore shed light on the validity of several taxonomic groups that have been erected among the Macrostomidae solely based on morphological synapomorphies (e.g. An-der-Lan 1939; Ferguson 1954; Faubel et al. 1994). Specifically, recent work suggests that there is widespread convergent evolution of numerous reproductive morphology traits, including the stylet (male intromittent organ), the sperm, the female antrum (female sperm receiving organ) (Schärer et al. 2011), as well as the presence of a second female opening (Schärer et al. 2011; Xin et al. 2019), possibly rendering several of the proposed synapomorphies paraphyletic.

In the current study, we present results from a global sampling effort to increase both taxon sampling and geographic representation of the genus. We also present a phylogenomic analysis based on a combination of transcriptome and Sanger-sequencing, thereby significantly expanding the phylogenetic and genomic resources available for the genus.

## 2. Material and Methods

### 2.1. Included species and documentation

In a global sampling campaign, we collected >1600 specimens of 131 *Macrostomum* species (see species delimitation section below) and additionally included data on 14 species from the literature. Almost all specimens were collected from the field in freshwater, brackish or marine habitats, either from sediment samples or from water plants (for details on specimens see Table S1). We documented specimens extensively with digital photomicrography, as previously described (Ladurner et al. 2005; Schärer et al. 2011; Janssen et al. 2015), using light microscopes (Leica DM2500, Olympus BH2, Leitz Diaplan, Zeiss Axioscope 5) with differential interference contrast (DIC) and equipped with digital cameras (Sony DFW-X700, Sony DFK41 and Ximea xiQ MQ042CG-CM). We collected both images and videos at various magnifications (40x to 1000x), documenting the general habitus and details of internal organs, which is possible due to the small size and high transparency of these organisms. To document sperm morphology, we amputated a worm’s tail slightly anterior of the seminal vesicle, ruptured the seminal vesicle (as described in Janicke and Schärer 2010), and documented the released sperm in a smash preparation using DIC and sometimes also phase-contrast microscopy. When possible, we prepared whole-mount permanent preparations of these amputated tails to preserve a physical specimen of the male intromittent organ, the stylet (Schärer et al. 2011; Janssen et al. 2015). Finally, we preserved the entire animal, or its anterior portion when amputated, for molecular analysis, in either RNAlater solution (Sigma, stored at 4°C up to a few weeks and then at −80°C) or in absolute ethanol (stored cool for up to a few weeks and then at −20°C).

### 2.2. Sequence data generation

We extracted both DNA and RNA from the RNAlater samples using the Nucleospin XS Kit in combination with the Nucleospin RNA/DNA Buffer Set (Macherey-Nagel), and we extracted DNA from the ethanol samples using the DNeasy Blood and Tissue kit (Qiagen, Germany). Extracted DNA and RNA was stored at −80°C. We amplified a partial *28S rRNA* sequence from DNA samples using PCR primers ZX-1 and 1500R and, for some fragments, additional nested PCR using primers ZX-1 and 1200R, or 300F and 1500R, with polymerases and cycling conditions as previously described (Schärer et al. 2011, 2020). We sequenced the resulting fragments from both sides using the PCR primers (Microsynth, Switzerland), assembled them in Geneious (v 11.1.5) with the built-in assembler (using the Highest sensitivity setting), with minor manual trimming and correction. For some fragments, we obtained additional sequences using internal primers 1090F, ECD2 (both Schärer et al. 2011).

To generate the RNA-Seq library for *M. clavituba* we extracted RNA from 40 pooled animals using Tri™ reagent (Sigma) and then prepared the library using the TruSeq® Stranded mRNA kit (Illumina). For all other selected RNA samples, we generated RNA-Seq libraries using SMART-Seq v4 (Clontech Laboratories, Inc.) combined with the Nextera XT DNA library preparation kit (Illumina). When at least 5ng total RNA was available, we performed the SMART-Seq v4 protocol with 12 preamplification steps and otherwise used 1ng and performed 15 preamplification steps. Libraries were checked for quality using a Fragment Analyzer (Agilent) and then sequenced as 101 paired-end reads on the HiSeq2500 platform (using the HiSeq® SBS Kit v4, Illumina) at the Genomics Facility Basel of the University of Basel and the Department of Biosystems Science and Engineering of the ETH Zürich.

### 2.3. Species delimitation

Most collected specimens could not be assigned to a taxonomically described *Macrostomum* species (http://turbellaria.umaine.edu, Tyler et al. 2006) and are likely new to science. We present transcriptome-level information for most species, but it was not feasible to conduct RNA-Seq on all >1600 collected specimens. We, therefore, relied on morphology and the partial *28S rRNA* sequences for species assignment since this fragment is widely used as a DNA barcode for flatworms (e.g. Chambrier et al. 2004, 2015; Janssen et al. 2015; Scarpa et al. 2015). We constructed haplotype networks from the partial *28S rRNA* sequences using the TCS algorithm (Templeton et al. 1992) implemented in the TCS software (Clement et al. 2000) and chose to delimit species with >3 mutational differences in the network. Such a difference in this *28S rRNA* fragment indicates distinct species among microturbellarians (Scarpa et al. 2015), but it is insufficient to clearly distinguish recently diverged species within *Macrostomum* (Schärer et al. 2020). Additional markers like mitochondrial COI are frequently used to detect recent divergence (e.g. Schärer et al. 2011; Janssen et al. 2015), but despite considerable efforts, we were unable to develop universal primers (a common issue in flatworms, see e.g. Vanhove et al. 2013) and individual primer optimization, as in Schärer et al. (2020), was not feasible here. We, therefore, chose to also delimit species with ≤3 differences in this *28S rRNA* fragment, provided that they showed clear diagnostic differences in morphology. We thus err on the side of lumping specimens, with species with shallow molecular and no diagnostic morphological divergence being assigned to the same operational species. We provide haplotype networks accompanied by drawings of the diagnostic features for all species in the supporting information (SI Species delimitation). Specifically, when a species’ morphology was diagnostic, we sequenced several specimens, if possible from different sample locations, to confirm that they were indeed molecularly similar and then assigned additional specimens based on morphology only. When no specific diagnostic traits could be defined (as was the case for many species of the hypodermic clade), we sequenced all specimens collected for molecular assignment. We used 668 sequences, of which 604 were generated for this study (Accessions: MT428556-MT429159) and 64 were from public databases (see also Table S1), representing the available diversity across the genus. We also included 16 sequences from seven species of our chosen outgroup genus *Psammomacrostomum* (see Table S1). We aligned sequences using MAFFT (“-genafpair -maxiterate 1000”) and generated haplotype networks for the recovered clades. We removed all columns that contained at least one gap before running TCS, leaving us with an alignment with 787 sites and 385 variable bases. We thus ignore indels to avoid the generation of cryptic species solely based on these, since scoring indels is non-trivial, and they are usually treated as missing data (Simmons et al. 2007).

### 2.4. Transcriptome selection

We used 134 transcriptomes representing 105 species, including four distant outgroups, *Haplopharynx rostratum, Microstomum lineare, Myozonaria bistylifera*, and *Karlingia* sp. 1, three species from the sister genus, *Psammomacrostomum* (see also Janssen et al. 2015), and 98 *Macrostomum* species in the phylogenomic analysis. This included the four publicly available high-quality transcriptomes of *M. hystrix, M. spirale, M. pusillum* (Brand et al. 2020), and *M. lignano* (Wudarski et al. 2017; Grudniewska et al. 2018), as well as 130 *de novo* assembled transcriptomes. Transcriptomes were assembled as previously described (Brand et al. 2020) and were mostly derived from whole single specimens and a few from anterior fragments only, which may reduce the transcriptome repertoire. Eight transcriptomes were assembled from combined single worm RNA-Seq data sets collected at the same location and assigned to the species based on our taxonomic expertise. Finally, the transcriptome of *M. clavituba* was assembled from RNA extracted from 40 pooled animals collected from the same location (for details see Table S2).

We assessed transcriptome quality using TransRate (version 1.0.2, Smith-Unna et al. 2016), which maps the reads back to the assembly and calculates mapping metrics, and BUSCO (version 2.0, Waterhouse et al. 2017, 978 genes in the metazoan dataset, version 2016-11-01), which searches for the presence of a curated set of core conserved genes. BUSCO scores were also used to select one representative transcriptome when multiple transcriptomes were available for a species (see below). We determined the empirical insert size of our libraries by mapping the reads to the assemblies using SNAP (version 1.0, Zaharia et al. 2011) and then extracting the mean insert size using Picard (version 2.20.2).

### 2.5. Orthology inference

To infer a set of orthologous genes (orthologs) we predicted open reading frames (ORFs) for each transcriptome using TransDecoder (version 5.3.0, Haas et al. 2013) with Pfam searches (version 32.0) to retain transcripts with predicted proteins and kept only one ORF per transcript using the “single_best_only” option. We then clustered predicted proteins with at least 99.5% sequence identity using the CD-HIT clustering algorithm (version 4.7, Fu et al. 2012). The amino acid sequences where then processed with OrthoFinder (version 2.2.6, Emms and Kelly 2015) using the “-os” option to perform length-adjusted reciprocal BLAST searches, followed by MCL clustering. We processed the resulting set of homologous genes (homogroups) using modified scripts from the phylogenomic dataset construction workflow (Yang and Smith 2014). We aligned all homogroups that contained at least 10 species using MAFFT (version 7.310, “--genafpair --maxiterate 1000”, Nakamura et al. 2018), inferred the best substitution model with ModelFinder (Kalyaanamoorthy et al. 2017), and the gene tree using IQ-TREE (version 1.5.5, Nguyen et al. 2015, command: “-mset DCMut, JTTDCMut, LG, mtZOA, PMB, VT, WAG -mfreq FU,F -mrate G”). Then we trimmed the gene trees using “trim_tips.py” to remove tip branches that were longer than two substitutions per site and “mask_tips_by_taxonID_transcripts.py” to remove monophyletic paralogs by choosing the sequence with the best representation in the alignment. We split off subtrees with long internal branches using “cut_long_internal_branches.py” and inferred orthologs using the rooted outgroup method in “prune_paralogs_RT.py”. This method uses known outgroup taxa to root the phylogenies, which then allows to infer the history of speciation and duplication and extract the most inclusive set of orthologs. We defined *Haplopharynx rostratum, Microstomum lineare, Myozonaria bistylifera*, and *Karlingia sp. 1* as the outgroup and all *Macrostomum* and *Psammomacrostomum* as an ingroup, since the algorithm does not include the defined outgroup in the output, and we then used *Psammomacrostomum* to root our final ortholog trees. Next, we realigned the orthogroups using MAFFT and trimmed the alignment with ZORRO (Wu et al. 2012), discarding any columns in the alignment with a score of less than five and filtering alignments that were shorter than 50 amino acids after trimming. Finally, we inferred ortholog gene trees with 100 non-parametric bootstraps with IQ-TREE, by inferring the best fitting model (“-mset DCMut, JTTDCMut, LG, mtZOA, PMB, VT, WAG -mfreq FU,F -mrate E,I,G,I+G”). These best fitting substitution models were later also used for the partitioned maximum-likelihood analysis. We generated two gene matrices, one matrix containing many genes but a relatively moderate occupancy (called L for low occupancy) and one with a lower gene number but a higher occupancy (called H for high occupancy, Table 1).

**Table 1.** Characteristics of the protein alignments used for phylogenetic inference. We used one alignment aimed at a high number of genes (L alignment) and one alignment aimed at high occupancy (H alignment). The alignments used were trimmed to only include regions with a high probability of being homologous using ZORRO, and the statistics are given for the alignments before and after trimming.

| Alignment metrics | L untrimmed | L | H untrimmed | H |
| --- | --- | --- | --- | --- |
| No. genes | 8128 | 8128 | 385 | 385 |
| No. amino acids AA | 5,057,157 | 1,687,014 | 200,729 | 94,625 |
| No. variable sites | 3,263,955 | 1,157,689 | 135,887 | 74,175 |
| No. parsimony informative sites | 2,287,246 | 934,803 | 103,425 | 63,066 |
| Missing data (%) | 78.1 | 59.3 | 55.7 | 22.9 |

### 2.6. Phylogenetics

We conducted most analyses on the H alignment for computational tractability and since missing data can have a negative influence on tree inference, particularly when using Bayesian methods (Roure et al. 2013, but see Tan et al. 2015), and only the less computationally demanding summary and maximum-likelihood methods were also applied to the L alignment. So for both alignments, we calculated a maximum likelihood species phylogeny using IQ-TREE (“L-IQ-TREE” and “H-IQ-TREE”) with a partition for each gene and using the best substitution model for each gene (see above). We calculated 1000 ultrafast bootstraps and conducted an approximate-likelihood ratio test to assess branch support.

Additionally, we inferred Bayesian phylogenies using ExaBayes (version 1.5, Aberer et al. 2014) for the H alignment (“H-ExaBayes”) with partitions for each gene, equal prior probability on all available amino acid substitution models, and with gamma models for all partitions. We ran four independent chains retaining every 500th iteration and discarding the first 365,000 iterations as burn-in. We terminated the analysis after 1.46 million generations since the average deviation of split frequencies between all the chains was below 1%, indicating convergence. We further assessed convergence using Gelman’s R of the likelihoods with the R package coda (Plummer et al. 2006), which showed three chains had converged, while one appeared stuck on a local peak. Since all chains quickly converged on the same topology, the local peak probably occurs due to differences in the substitution models applied, with the HIVB model being present at a low rate in the three converged chains, but being absent in the fourth. We combined the three converged chains, discarded the fourth and calculated a consensus tree using quadratic path distance optimization (using the ls.consensus function in the R package phytools).

To account for potential issues caused by model misspecification, we also performed an unpartitioned analysis using the CAT model implemented in PhyloBayes (version 1.5, Lartillot et al. 2013). Because a full GTR model of the amino acids was too parameter rich, we ran the tool on the DNA coding sequence of the H alignment and fit the CAT+GTR model (“H-PhyloBayes”). Two chains were run in parallel on 400 CPUs for two weeks. Due to the high cost of the analysis, we terminated the run after 23,311 iterations, at which point the chains showed an identical topology (after removal of 10,000 iterations as a burn-in). At this point the realized difference in the likelihoods between the two H-PhyloBayes chains was high (0.21 with an ESS of 858) suggesting the chains had not converged. However, Gelman diagnostics for the likelihoods suggested convergence. Our chains could be at a local peak where the hypodermic clade is paraphyletic, possibly due to poor mixing, potentially biasing the results. Therefore, we discuss the H-PhyloBayes phylogeny, but exclude it from the comparative analysis.

To construct phylogenies while accounting for potential incomplete lineage sorting, we ran the quartet based method implemented in ASTRAL-III (version 5.6.1, Zhang et al. 2018) on both alignments (“L-ASTRAL” and “H-ASTRAL”). We assessed the level of gene tree – species tree conflict at each node of the phylogeny using the quartet support score, which gives the proportion of quartet trees induced by the gene trees that are supporting the species tree topology as opposed to the two possible alternatives (Zhang et al. 2018). Strong support for the species tree partition can be interpreted as little disagreement between the gene trees, while support for the alternate topologies indicates strong gene tree – species tree conflict.

Furthermore, we chose representative *28S rRNA* sequences from the haplotype network analysis to infer phylogenetic placement for species without a transcriptome. For species with a transcriptome, we mostly chose the *28S rRNA* sequence derived from the same specimen (for exceptions see Table S2). We aligned the sequences using MAFFT (“--maxiterate 1000 --globalpair”), trimmed the start and end using trimAl (“--nogaps --terminalonly”) (Capella-Gutierrez et al. 2009) and determined the best substitution model using ModelFinder with the BIC criterion. We then combined this alignment with the H alignment, and analysed it with the best fitting substitution model using IQ-TREE as described above (referred to as “C-IQ-TREE”, called C for combined). To facilitate comparative analysis, we transformed the phylogenies (C-IQ-TREE, H-IQ-TREE and H-ExaBayes) to be ultrametric and with a root depth of 1 using the penalized marginal likelihood approach (Sanderson 2002) implemented in the software TreePL (Smith and O’Meara 2012). After all of the above analyses were completed, we discovered that some transcripts of the RNA-Seq libraries constructed with the SMART-Seq v4 cDNA kit contained cDNA synthesis primer sequences. We did not remove these primers before transcriptome assembly since i) the manufacturer specifically states they should not occur in RNA-Seq reads if used in combination with the Nextera XT DNA library preparation kit (Appendix C in SMART-Seq v4 Ultra Low Input RNA Kit for Sequencing User Manual, Takara Bio Inc. available at: https://www.takarabio.com/assets/a/114825) and ii) because the relevant primer sequence is proprietary (both confirmed by Takara EU tech support in November 2020), so that we initially did not have access to its sequence. We only discovered the offending primer sequence due to our work on two ongoing *Macrostomum* genome projects. Because the analyses presented here had at that point used approximately 650,000 CPU hours of computation, and since we expected the effect of that primer to be small, we elected to perform follow-up analyses to determine how robust our results were to its removal, rather than opting for a complete re-assembly and re-analysis of all the data. As we outline in the supporting information (SI Primer removal), the removal of the cDNA synthesis primer sequences had little effect on the topology and branch lengths of L-IQ-TREE, H-IQ-TREE and C-IQ-TREE and we, therefore, consider our results to be robust.

## 3. Results and Discussion

### 3.1. Phylogenetics

#### 3.1.1. Large undescribed species diversity

Based on an integrative approach—using detailed documentation of *in vivo* morphology and *28S rRNA* sequences—we identified 51 species as previously described and a striking 94 species as likely new to science (Fig. 2, Table 2), thereby increasing the species diversity in the genus by about 50% (http://turbellaria.umaine.edu). To facilitate future taxonomic work and comparative analysis of this diversity, we have deposited extensive image and video data (see Brand and Schärer 2021 for a Zenodo deposit), a vector drawing file of Fig. 2 (Fig. S1, which includes our stylet and sperm drawings), as well as geographic and molecular data for all the specimens collected (see Table S1 and S2; with each specimen carrying a unique Macrostomorpha Taxonomy and Phylogeny ID number taking the form of “MTP LS ##”; see also http://macrostomorpha.info). Moreover, we assigned specimens for which we do not have molecular data to operational taxonomic units based on morphology.

**Table 2.** Summary of the taxonomic status of all the included *Macrostomum* species. Species inferred based on single specimens (N=1) are listed separately.

| Species in this study | N = 1 | N > 1 | Total |
| --- | --- | --- | --- |
| All | 18 | 127 | 145 |
| Described | 5 | 46 | 51 |
| Likely undescribed & immature | 3 | 2 | 5 |
| Likely undescribed, mature & no transcriptome | 5 | 17 | 22 |
| Likely undescribed, mature & transcriptome | 5 | 62 | 67 |

#### 3.1.2. Phylogenetic inference is robust to the methods applied

Across the different phylogenies, all species represented by more than one transcriptome (i.e. 15 x 2, 3 x 3, and 1 x 9 transcriptomes) were monophyletic (Fig. S2), with the notable exception of *M. lignano* (see below). The grouping of the major clades was mostly consistent across all phylogenetic approaches (Fig. 3 and Fig. S2). All phylogenies recovered seven large species groups (further called the hypodermic [further divided into hypodermic I and II], spirale, lignano, finlandense, tuba, and tanganyika clade), two smaller species groups (the minutum and hamatum clade), and two consistent species pairs (*M*. sp. 45 + 46 and *M*. sp. 4 + 89). However, the backbone, and the positions of some species with long branches (*M*. sp. 37, *M*. sp. 39, *M*. sp. 90 and *M. curvituba)*, differed depending on the alignment and method used. Despite these discrepancies, the Robinson-Foulds distances (Robinson and Foulds 1981) between the phylogenies were low (Table S3), indicating good agreement between the methods.

**Fig. 3.**
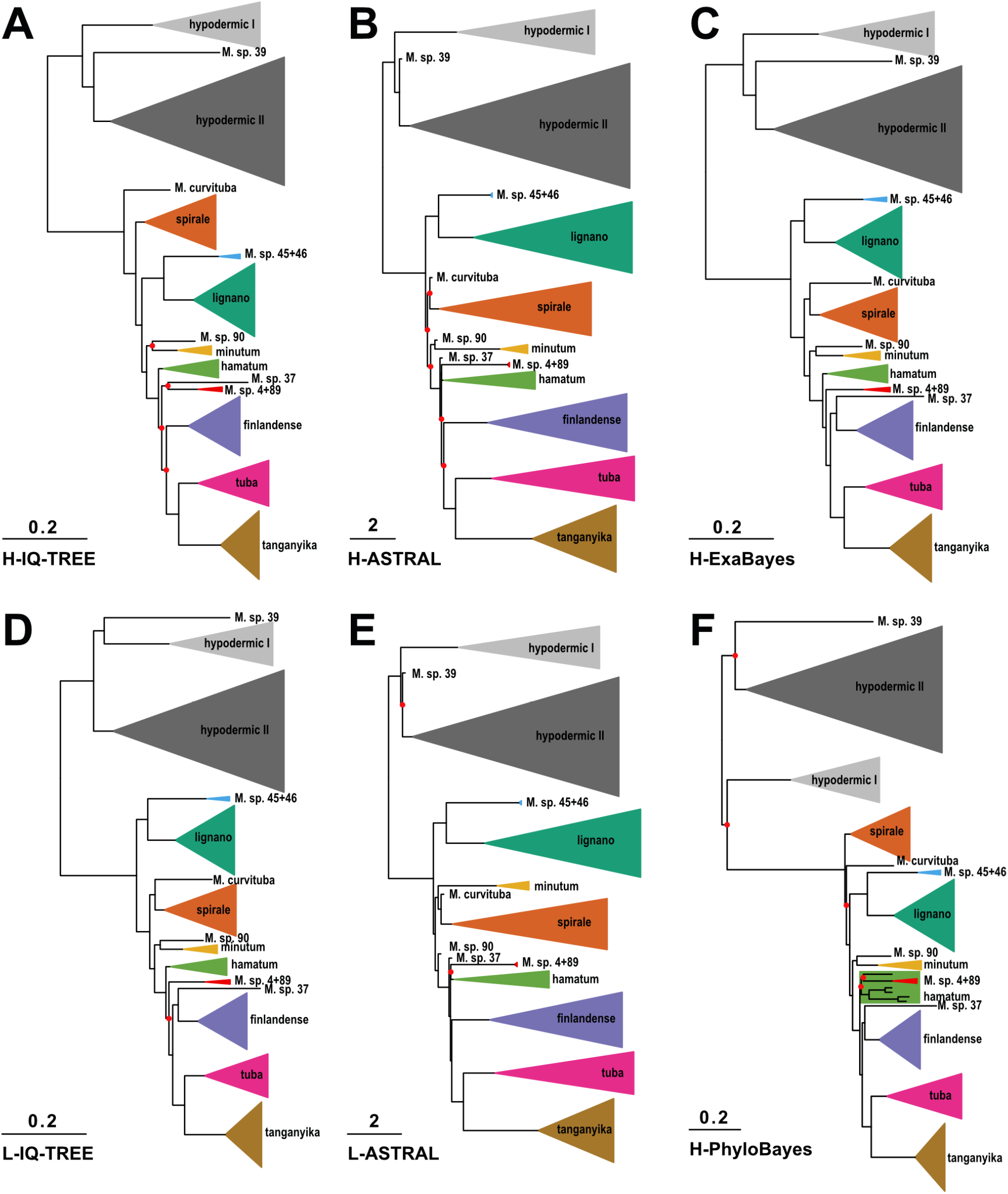
Simplified transcriptome-based phylogenies of the genus *Macrostomum* inferred using various methods on the L and H alignments (see also Table 1). Consistently recovered clades are collapsed into named triangles, with the height of the base proportional to the number of species, and single species names preceded by the genus abbreviation. Clades are coloured consistently between the panels, with seven large species groups (hypodermic I, light grey; hypodermic II, dark grey; spirale, orange; lignano, dark green; finlandense, purple; tuba, pink; tanganyika, brown), two smaller species groups (minutum, yellow; hamatum, light green), and two consistent species pairs (*M*. sp. 45 + 46, light blue; *M*. sp. 4 + 89, red). Branch length represents substitutions per site in A, C, D and F and coalescent units in B and E. Bipartitions that did not have maximal support are indicated with red circles (for details see Fig. S2).

In all phylogenies, the hypodermic clade was deeply split from the reciprocal clade, but with large phylogenetic distances also appearing within the hypodermic clade (referred to as hypodermic I and II) (Fig. 3 and Fig. S2). In H-PhyloBayes the hypodermic clade was not monophyletic, but we did not find maximal node support for these splits, and since we had issues with the convergence of the PhyloBayes runs, we are skeptical of this alternative topology. Within the reciprocal clade the tanganyika, tuba and finlandense clades were consistently grouped and the hamatum and minutum clades, together with *M*. sp. 4 + 89, were always the closest relatives to the former three clades. However, the exact relatedness patterns were uncertain. In the L-IQ-TREE, H-IQ-TREE and H-ExaBayes phylogenies, *M*. sp. 4 + 89 were the closest relatives to these three clades, followed by the hamatum and minutum clades. In contrast, in the ASTRAL and the H-PhyloBayes phylogenies, the hamatum clade was more closely related to the minutum clade (with the latter nested within the former in case of H-PhyloBayes) and both were sister to the grouping of tanganyika, tuba and finlandense. Moreover, the exact branching order at the base of the reciprocal clade was not clearly resolved. The spirale clade split off first in the H-IQ-TREE and H-PhyloBayes phylogenies, while the lignano clade split off first in the other phylogenies.

Consistent with the conflict between methods, the quartet support from the ASTRAL analysis indicated gene tree – species tree conflict at most nodes in the phylogeny’s backbone (Fig. S3). These internal nodes were separated by short branches suggestive of rapid speciation events, such as during adaptive radiation (Irisarri et al. 2018), where substantial incomplete lineage sorting is expected. This pattern is also consistent with ancient hybridization, which is a distinct possibility, since there is evidence for hybridization between the sibling species *M. lignano* from the Eastern Mediterranian and *M. janickei* from the Western Mediterranian under laboratory conditions (Singh et al. 2020).

In this context, and with respect to the above-mentioned population divergence within *M. lignano*, it is interesting to point out that one *M. lignano* transcriptome was from the DV1 inbred line from the type locality near Bibione, Northern Adriatic Sea, Italy (Vellnow et al. 2017), while the other was from an outbred population from the Sithonia Peninsula, Aegean Sia, Greece (Zadesenets et al. 2016; Schärer et al. 2020). *M. lignano* was monophyletic in the L-IQ-TREE, H-PhyloBayes, and both ASTRAL phylogenies, but the Greek population was sister to *M. janickei*, from near Montpellier, France, in the H-IQ-TREE and H-ExaBayes phylogenies, although node support for this split was low for H-IQ-TREE (see dark green clade in Fig. S2). Moreover, removal of cDNA synthesis primer sequences from the alignment rendered *M. lignano* monophyletic also in H-IQ-TREE (see SI Primer removal, Fig. A4). A closer investigation of the morphology of the Greek vs. Italian population also revealed that the former had a considerably larger stylet (96.7 vs. 60.7 µm) and longer sperm (76.5 vs. 62.7 µm) (data not shown). So while we still consider *M. lignano* a proper species here, a closer comparison of these Greek and Italian populations would be interesting, as well as a broader sampling of *M. janickei*.

The topology of C-IQ-TREE (Fig. 2) was identical to H-IQ-TREE (Fig. 3A and Fig. S2A) when we removed all the species added based on only *28S rRNA* and thus adding these species did not negatively influence the overall topology of the tree. Node support in C-IQ-TREE was somewhat lower, as could be expected given the placement of the additional species based solely on *28S rRNA* sequences. Nevertheless, we think that a ∼50% increase in species representation is highly worthwhile, and we focus on this combined C-IQ-TREE phylogeny in the following.

#### 3.1.3. Cryptic diversity in the hypodermic clade

As already mentioned above, many species in the hypodermic clade were highly molecularly diverged, even though they are difficult to distinguish morphologically. Most of these species had stylets that consisted of a short proximal funnel that tapered to a curved and drawn-out asymmetrical needle-like thickening (Fig. 2A, top). It was possible to distinguish some species based on general habitus, and these differences are reflected by the four deeply split clades containing, *M. rubrocinctum, M. hystricinum, M. gabriellae*, and *M. pusillum*, respectively (although the latter two clades were also quite similar in habitus). We have indicated substantial morphological similarity by appending a letter to the species name (e.g. *M. hystricinum a, c, d*).

Investigations of species within the hypodermic clade without support from molecular data thus require considerable caution, and earlier authors have already identified considerable difficulties in distinguishing species with this morphology (e.g. Beklemischev 1951; Luther 1960; Rieger 1977; Ladurner et al. 1997), which is even more poignant due the considerable levels of convergent evolution in the genus resulting from evolutionary shifts to hypodermic insemination (Schärer et al. 2011), which we analyse in more detail in a separate study (Brand et al. 2021). Moreover, given these striking morphological similarities, it may often not be clear to which of these species the name-bearing type specimens belong to (Schärer et al. 2011), so the species names in this clade should be considered tentative. Eventually, one may need to name these species afresh, applying more extensive molecular species delimitation, and either suppress the original names of species without detailed enough morphological descriptions and/or lacking adequate type material, or to define neotypes, including molecular voucher specimens (Pleijel et al. 2008).

### 3.2. Morphological diversity

#### 3.2.1. A novel stylet morphology in the hypodermic clade

While the hypodermic clade showed little variation in terms of stylet and sperm morphology, the stylet of *M*. sp. 93 (Fig. 2A top) differed clearly from that stereotypical form, by having a small proximal funnel extending into a straight and obliquely-cut tube (Fig. 4A). This shape is similar to the stylets of several species in the reciprocal clade, namely *M. shenda, M*. sp. 34 (both Fig. 2A bottom) and *M*. sp. 64 (Fig. 2B bottom), as well as *M. orthostylum* (for which we have no phylogenetic placement). As we will report in more detail in a separate study (Brand et al. 2021), we observed hypodermic received sperm in both *M*. sp. 93 and *M*. sp. 64, and this stylet shape thus appears adapted for hypodermic insemination. While we did not observe hypodermic received sperm in *M. orthostylum* or *M*. sp. 34 (nor was such sperm reported for *M. shenda* by Xin et al. 2019), these species are nevertheless likely to also mate through hypodermic insemination, not least since stylets with similar shapes are also used for hypodermic insemination in related macrostomid flatworms (Janssen et al. 2015).

**Fig. 4.**
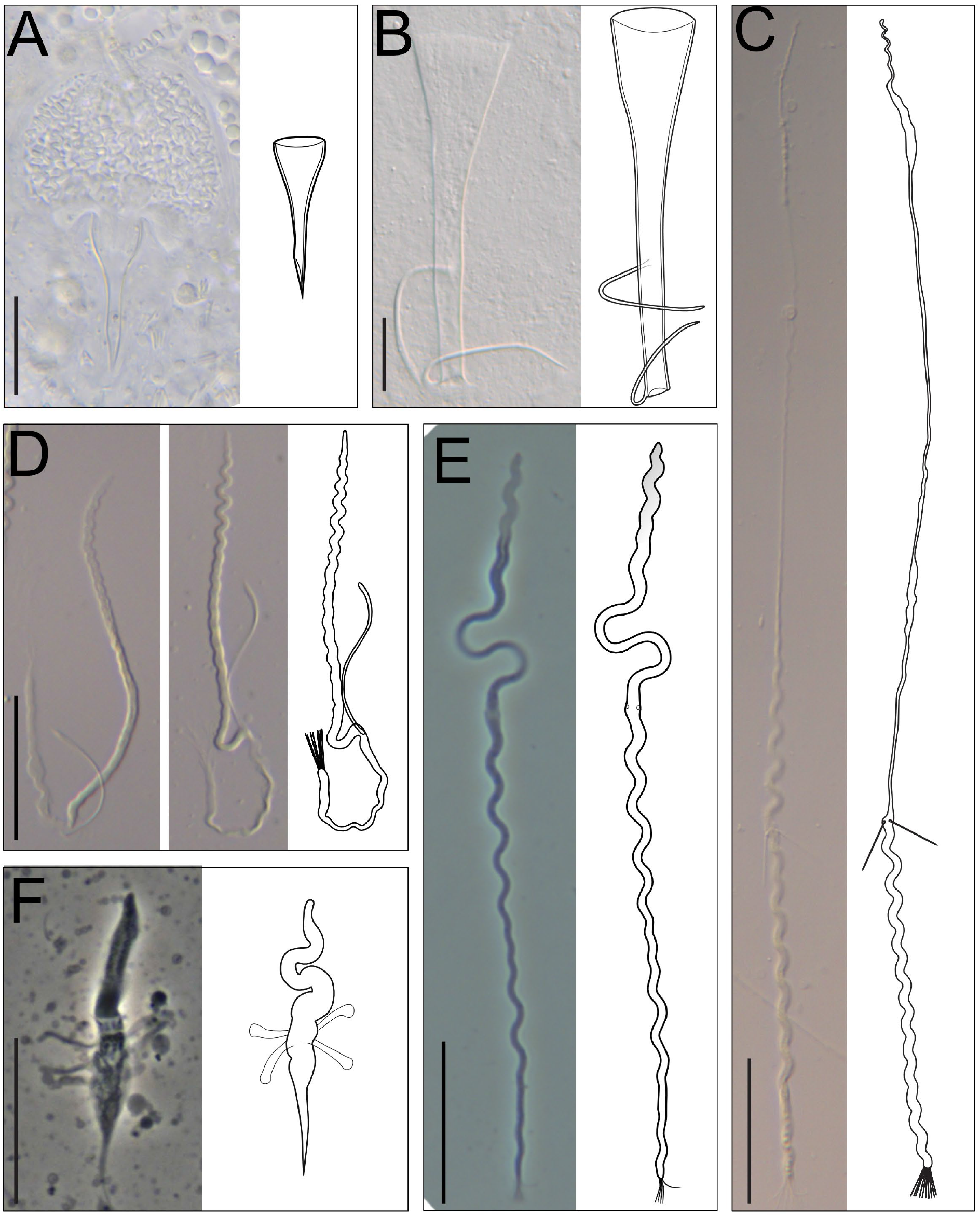
Phase contrast or DIC micrographs and line drawings of novel stylet (A-B) and sperm (C-F) morphologies. The brackets give the specimen ID of the micrographs. (A) The stylet of *M*. sp. 93 is atypical for a representative of the hypodermic clade (MTP LS 2985, DIC). (B) The stylet of *M*. sp. 43 has two thin lateral protrusions (MTP LS 2106, DIC smash preparation). (C) The elongated sperm of *M*. sp. 67 (MTP LS 2613, stitched from three images, DIC). (D) Two micrographs showing the sperm of *M*. sp. 42 with only a single, flexible sperm bristle (MTP LS 3215, DIC). (E) The sperm of *M*. sp. 82 with an anterior region that appears translucent (MTP LS 2877, phase contrast). (F) The sperm of *M. distinguendum* with several lateral appendages (MTP LS 602, phase contrast). Scale bars represent 20 µm.

#### 3.2.2. Diversification of stylet and sperm morphology

In contrast to the largely canalized stylet and sperm morphology of the hypodermic clade, we found remarkable variation in these structures within the reciprocal clade (Fig. 2). For example, we documented five species in the spirale clade that had stylets with lateral protrusions close to the distal opening (Fig. 2A middle). The protrusion is shaped like a rod or spike in *M. evelinae, M*. sp. 29 and *M*. sp. 42, while it consists of two thin filaments in *M*. sp. 30 and *M*. sp. 43 (Fig. 4B; see also Fig. A13 in SI Species delimitation).

These highly modified stylets may have coevolved with the complex antra in these species, which have two separate chambers connected via a ciliated muscular sphincter. The sperm of these species is also remarkable because they carry only a single bristle instead of the typical two. The sperm of *M*. sp. 30 carry only a single curved bristle, while the bristle of *M. evelinae, M*. sp. 29, and *M*. sp. 42 (Fig. 4D) is additionally modified, being thicker and appearing flexible, frequently curving back towards the sperm body. A second bristle might also be absent in the closely related *M*. sp. 13 (indicated with a shaded second bristle in Fig. 2A middle), but the available material currently does not allow an unambiguous assessment.

Sperm are also highly variable across the entire reciprocal clade. Particularly striking are the sperm modifications of *M*. sp. 82 (Fig. 2B top), which give the anterior part of the sperm a translucent appearance under phase contrast (Fig. 4E), and which provides motility that is different from that observed in normal sperm feelers. We also document extraordinarily long sperm in several species in the tanganyika clade (Fig. 2B bottom). While it is not entirely clear which part of the sperm is modified here, it appears that they have very long sperm feelers, with the bristles thus being located unusually far posterior, as in *M*. sp. 67 (Fig. 4C). These long feelers represent a novel character in this African clade and should be interesting targets of future investigations. Additionally, we observed numerous species that had short or no sperm bristles across the whole reciprocal clade and appeared to coincide with changes in the stylet morphology. But as already mentioned above, a more formal analysis of convergent evolution in stylet and sperm morphology, including such reductions and losses of sperm bristles, will be presented in a separate study (Brand et al. 2021). Finally, a striking modification of sperm design occurs in *M. distinguendum* in the finlandense clade (Fig. 2B middle), which appears to lack sperm bristles, but instead carries novel club-shaped lateral appendages, which in light microscopic images bear no resemblance to the usual sperm bristles (Fig. 4F).

### 3.3. Taxonomic implications

The large-scale collections and molecular phylogenetic analyses we present here have a number of implications for the taxonomy and nomenclature of the Macrostomidae (Tyler et al. 2006), and they support several taxonomic and nomenclatural revisions. We outline these changes in the following, using the “*Genus species* Author, Year” format to refer to binomials, and additionally including all the citations for the relevant works.

#### 3.3.1. *Bradburia* is questionable

As part of the description of some Australian macrostomids, Faubel et al. (1994) established the genus *Bradburia* Faubel, Blome & Cannon, 1994 for species with a “Male system with accessory stylet connected to tubular stylet proximally” as the most prominent diagnostic trait. They named a new species, *Bradburia australiensis* Faubel, Blome & Cannon, 1994, and transferred a species originally described as *Macrostomum miraculicis* Schmidt & Sopott-Ehlers, 1976 (Schmidt and Sopott-Ehlers 1976) to this genus, designating it *Bradburia miraculicis* (Schmidt & Sopott-Ehlers, 1976) and making it the type species of the genus *Bradburia*.

While we have collected neither of these two species ourselves, we have identified several species that carry some kind of lateral protrusion on the stylet shaft. These include *Macrostomum inductum* Kolasa, 1971 (Kolasa 1971) and *Macrostomum* sp. 108 (Fig. 2B middle and Fig. 2A bottom, respectively), which have independently evolved ridge-like protrusions on the distal stylet. In addition, we have recovered a clade that includes *Macrostomum evelinae* Marcus, 1946 (Marcus 1946), in which all but one species carries elongated lateral protrusions on the stylet shaft, albeit not proximally, but rather in the distal third of the stylet (Fig. 2A middle). Note that also in *B. miraculicis* the lateral protrusion is not actually positioned proximally on the stylet—as the diagnosis of the genus *Bradburia* would imply—but according to Schmidt & Sopott-Ehlers (Schmidt and Sopott-Ehlers 1976) it emerges about half-way (“Etwa auf halber Länge …”). Moreover, Schmidt & Sopott-Ehlers draw that lateral protrusion as an integral part of the stylet, rather than an accessory stylet (as is, for example, observed in the Dolichomacrostomids), which therefore does not conform to the *Bradburia* genus diagnosis. In contrast, the lateral protrusion of the stylet of *B. australiensis*, as drawn by Faubel et al., could possibly be somewhat more independent.

Other diagnostic traits of the genus *Bradburia* are also not decisive, including the absence of eyes (a widespread trait in the genus *Macrostomum*, including the above-mentioned *M. evelinae* clade) and the presence of cuticularised parts in the female antrum, which are, for example, also observed in *Macrostomum ermini* Ax, 1959 (Fig. 2A middle) (Ax 1959, a species that lacks lateral stylet protrusions). Surprisingly, the organization of the female system was not observed in the description of *B. australiensis*, which raises the question why Faubel et al. (1994) consider it *Bradburia*.

The above considerations suggest that the genus *Bradburia* is currently not well-supported and we think it likely that molecular analyses would lead to the placement of both *B. miraculicis* and *B. australiensis* in the reciprocal clade of the genus *Macrostomum*, and possibly even into the *M. evelinae* clade, which may thus lead to the eventual suppression of the genus *Bradburia*.

#### 3.3.2. *Promacrostomum* should be dropped

Based on specimens collected from Lake Ohrid, An-der-Lan (1939) established a new genus *Promacrostomum* An-der-Lan, 1939, with initially a single representative, *Promacrostomum paradoxum* An-der-Lan, 1939 based on the presence of two female genital openings and a genito-intestinal duct. In all our phylogenies An-der-Lan’s species was placed within the reciprocal clade as sister to *M. retortum* Papi, 1951 (Papi 1951). Moreover, a second female opening has independently evolved multiple times in the genus *Macrostomum* (Schärer et al. 2011; Xin et al. 2019), so this trait alone is clearly not a useful synapomorphy for this genus. We, therefore, transfer this species from *Promacrostomum* to *Macrostomum paradoxum* (An-der-Lan, 1939). Moreover, as outlined by Schärer et al. (2011), another species with two female genital openings, *Macrostomum gieysztori* Ferguson, 1939 (Ferguson 1939), also belongs to *Macrostomum*, despite it having previously been placed into *Promacrostomum* by Papi (Papi 1950, 1951) and later the genus *Axia* by Ferguson (1954).

With this reclassification, the genus *Promacrostomum* loses the type species and would only contain *Promacrostomum palum* Sluys, 1986 (Sluys 1986), which was assigned to this genus because it, like *M. paradoxum*, has two female genital openings (while a genito-intestinal duct was not observed by Sluys). While it seems likely that a molecular placement of this species would reveal that it belongs to *Macrostomum*, it is unclear whether it would group with any of the known representatives with this trait state or represent yet an additional independent origin. Given that the main characteristic of the genus *Promacrostomum*, namely the presence of two female genital openings, has several independent origins, we consider *Promacrostomum* unjustified and transfer its last remaining species from *Promacrostomum* to *Macrostomum palum* (Sluys, 1986).

#### 3.3.3. *M. zhaoqingensis* could be a re-description of *M. inductum*

The comparison of *28S rRNA* sequences has revealed that *Macrostomum zhaoqingensis* Lin & Wang, 2017 (Lin et al. 2017b) is sister to *Macrostomum inductum* Kolasa, 1971 (Kolasa 1971) and the haplotype analysis closely clusters both species (see Fig. A19 in SI Species delimitation), suggesting that *M. zhaoqingensis* could potentially represent a re-description of *M. inductum*. More detailed molecular and morphological comparisons are called for to determine the taxonomic status of *M. zhaoqingensis*.

### 3.4. Conclusions

Our detailed phylogenomic investigations of 145 *Macrostomum* species show that the genus consists of two well-separated clades. Across phylogenetic methods the grouping within the first, morphologically canalized, hypodermic clade is fairly robust, but the exact interrelationships within the, morphologically diverse, reciprocal clade are somewhat less clear. However, we also find highly consistent subclades within the reciprocal clade, with most conflicts between methods stemming from uncertainty in the backbone of the phylogeny. Short internal branches of all phylogenies, and low split support from the ASTRAL analysis, suggest that this likely occurs because of a rapid radiation at the base of the reciprocal clade. Such a radiation would then lead to substantial gene tree – species tree conflict due to incomplete lineage sorting.

Remarkably, 94 of the collected species are likely new to science, highlighting that a large proportion of the diversity in this genus is yet to be discovered. Our increased taxon sampling has not only yielded many more species, but these additional species have also revealed a range of novel morphological traits. While some of these traits are phylogenetically clustered, others, which have previously been considered diagnostic for the erection of separate genera, are paraphyletic, thus requiring several taxonomic changes. The striking convergence of a range of traits makes taxonomic assignments solely based on morphology questionable and we suggest that future work, particularly on species exhibiting the hypodermic morphology, should always include molecular markers. The need for employing molecular markers is further supported by the large cryptic diversity within the hypodermic clade, a taxonomically challenging group that, however, could be interesting for investigations of cryptic speciation.

## Acknowledgments

We thank the numerous people that have helped with field work. Especially, we are grateful for the help of, in no particular order, Werner Armonies, Benny Glasgow, Mohamed Charni, Edith Zemp, Bernhard Egger, Peter Ladurner, Gregor Schulte, Floriano Papi, Kazuya Kobayashi, Christopher Laumer, Wim Willems, Tom Artois, Christian Lott, Miriam Weber, Ana-Maria Leal-Zanchet, Kaja Wasik, Mariana Adami, Walter Salzburger, Adrian Indermaur, Bernd Egger, Fabrizia Ronco, Heinz Büscher, Victoria Huwiler, Philipp Kaufmann, Michaela Zwyer, Stefanie von Fumetti, Joe Ryan, Mark Q. Martindale, Marta Chiodin, John Evans, Leigh Simmons, Mauro Tognon, Piero Tognon, Cristiano Tognon, Pragya Singh, Nikolas Vellnow, Christian Felber, Ulf Jondelius, Sarah Atherton, Tim Janicke, Georgina Rivera-Ingraham, Ben Byrne, Yvonne Gilbert, Rod Watson, Jochen Rink, Miquel Vila-Fare, Helena Bilandžija, and Sasho Trajanovski. We thank Katja Eschbach of the Genomics Facility Basel for preparing and sequencing RNA-Seq libraries. We thank Peter Fields and Lukas Zimmermann for IT advice. We thank Jürgen Hottinger, Daniel Lüscher and Yasmin Picton for administrative and technical support. We thank Dita Vizoso for the use of sperm and stylet drawings for some of the previously described species and for providing inspiration for the new drawings. We thank Yu Zhang and his collaborators for kindly providing specimens of *M. baoanensis*. We thank the following locations and authorities for collection/export permits, Western Australia (DPAW Reg 17 01-000135-2 and Reg 18 OS002550), Victoria (DELWP National Parks Act 1975 10008144), Tenuta di San Rossore, Italy (3299/7-2-1/2010 and 1337/7-2-1/2016), Parco Nazionale Arcipelago Toscano, Italy (4440/2011), Sao Leopoldo, Brazil (IBAMA 12BR008859/DF), Carrie Bow Cay, Belize (STRI Field Station and Department of Fisheries), Lakes Ohrid and Prespa, North Macedonia (Hydrobiological Institute Ohrid), Swedish West Coast (Kristineberg Biological Station), Reserve Naturelle de la Petite Camargue Alsacienne, France (years 2016 and 2017), Florida, USA (Whitney Laboratory for Marine Bioscience), Sardinia, Italy (Regione Autonoma della Sardegna 0035419/2018), Zambia (based on a Memorandum of Understanding and KA/B/29/16), Malawi, (Museum Karonga and Fisheries Department).We thank Nick Goldman and Ziheng Yang for organizing the summer school on Computational Molecular Evolution, which has helped and inspired the first author to explore phylogenetics. Calculations were performed, in part, at sciCORE (http://scicore.unibas.ch/) scientific computing centre at the University of Basel.

## Data availability

The raw sequencing data generated for this study are available in the NCBI

Sequence Read Archive repository with the following accession: PRJNA635941. The partial *28S rRNA* sequence are available on NCBI with the following accessions: MT428556-MT429159. Extensive image and video material of all documented specimens are deposited on Zenodo at: 10.5281/zenodo.4482135. Transcriptome assemblies, gene alignments and phylogenetic trees are deposited on Zenodo at: 10.5281/zenodo.4543289.

## Author contribution

Jeremias N. Brand: Conceptualization, Data Curation, Formal Analysis, Investigation, Visualization, Writing – Original Draft Preparation, Writing – Review & Editing. Gudrun Viktorin: Investigation, Methodology. R. Axel W. Wiberg: Investigation, Writing – Review & Editing. Christian Beisel: Methodology, Resources. Lukas Schärer: Conceptualization, Data Curation, Funding Acquisition, Investigation, Project Administration, Resources, Supervision, Writing – Original Draft Preparation, Writing – Review & Editing

## 6. Supp. Figures

**Fig. S1. Vector file of Fig. 2 to facilitate reuse of our stylet and sperm drawings in future studies**.

see the file Fig_S1.svg

**Fig. S2. Detailed transcriptome-based phylogenies of the genus *Macrostomum* inferred using various methods on the L and H alignments**. Clades are coloured consistently between the panels, with seven large species groups (hypodermic I and II, dark grey; spirale, orange; lignano, dark green; finlandense, purple; tuba, pink; tanganyika, brown), two smaller species groups (minutum, yellow; hamatum, light green), and two consistent species pairs (*M*. sp. 45 + 46, light grey; *M*. sp. 4 + 89, dark blue). Transcriptomes that are placed differently between approaches are highlighted in red. Species assignments are given as a short six letter abbreviation in the name of the transcriptomes (three letters each for the genus and species, respectively). Branch length represents substitutions per site in A, C, D and F and coalescent units in B and E. Bipartition support values are indicated at internal nodes of the phylogenies and indicate ultrafast bootstrap and approximate-likelihood ratio test values in A and D, quartet support scores in B and E, and posterior probability in F. Posterior probability was 1 for all bipartitions in C.

see the file Fig_S2.pdf

**Fig. S3. The two ASTRAL phylogenies with quartet support shown for each node**. (A) L-ASTRAL phylogeny, and

(B) H-ASTRAL phylogeny. The pie charts at the nodes represent the quartet support for the three possible partitions at that node. Blue represents support for the species tree topology and orange and red represent support for one or the other of the two alternative partitions. Mostly blue pie charts indicate little conflict. This analysis indicates that there is substantial gene tree – species tree conflict throughout the genus.

see the file Fig_S3.pdf

## 7. Supp. Tables

**Table S1. Details on all specimens included in this study**. Includes information on species names, GenBank accessions, sampling dates and locations.

see the file Tab_S1.xlsx

**Table S2. Details on the transcriptomes used in this study**. Includes information on assembly names, specimens, library type, and library statistics.

see file Tab_S2.xlsx

**Table S3. Robinson-Foulds distance matrix between the various phylogenetic inference methods**. The first row gives the alignment used for inference (L or H) and the second row the inference software used.

see file Tab_S3.xlsx

