## Supporting information for "Large-scale phylogenomics of the genus *Macrostomum* (Platyhelminthes) reveals cryptic diversity and novel sexual traits"

#### SI Primer removal

As mentioned in the main text, we discovered that most of our transcriptome assemblies contained a cDNA synthesis primer sequence. Specifically, a 21nt sequence corresponding to a partial SMART-Seq v4 adapter (confirmed by Takara EU tech support in November 2020 and further called "short primer") could be reliably identified from the start of many assembled transcripts. That sequence is very similar to other SMART primers (e.g. the SMARTseq2 TSO primer, Picelli et al. 2013, and the SMARTer II A Oligo, Clontech Laboratories). This sequence probably corresponds to a fragment of both the "SMART-Seq v4 Oligonucleotide" and the "3' SMART-Seq CDS Primer II A" that are shown in figure 2 of the SMART-Seq v4 User manual (Appendix C in SMART-Seq v4 Ultra Low Input RNA Kit for Sequencing User Manual, Takara Bio Inc. available at: https://www.takarabio.com/assets/a/114825). The shorter length of this sequence compared to other SMART sequences is possibly due to incomplete cleavage of the primer by the Nextera XT protocol (suggested by Takara EU tech support in November 2020). We, therefore, inferred a longer 28nt sequence of a putative complete SMART-Seq v4 adapter and used it for trimming (further called "long primer", Fig. A1).

```
SMARTseq2 TSO primer    AAGCAGTGGTATCAACGCAGAGTACATrGrG+G
SMARTer II A Oligo      AAGCAGTGGTATCAACGCAGAGTACXXXXX
Short primer                   GGTATCAACGCAGAGTACGGG
Long primer             AAGCAGTGGTATCAACGCAGAGTACGGG
```

**Fig. A1. Known SMART primer sequences and the two sequences identified here.** The "short primer" sequence shares an 18nt region (blue) with two commonly used SMART primers, but is different in the terminal region (red). Since the proprietary primer used in the SMART-Seq v4 cDNA kit likely is longer than that sequence, we added the remaining common region (green) to it. Primer trimming efficiency increased slightly with the "long primer".

Across transcriptomes derived from libraries constructed using the SMART-Seq v4 cDNA kit, on average, 29.5% (range: 11.7% – 70.9%) of all contigs had a BLAST hit for the "short primer" sequence, and similarly, 10.4% (range: 4.3% – 29.4%) of the coding sequences resulting from TransDecoder protein predictions contained the "short primer" sequence. In contrast, none of the assemblies derived from other library construction techniques resulted in any BLAST hits (see Tab S3 for details on primer content for each assembly).

We next investigated if primer sequences were also present in the coding sequences of the final protein alignments. Alignment annotation using the Geneious (version 11.1.5) annotation tool showed that the primer was indeed present in the untrimmed alignments, but that it was mostly removed in the ZORRO alignment trimming step. Visual inspection suggested that, as expected, primers mostly occurred at the beginnings and ends of alignments and thus in regions often removed during the trimming process. However, in some cases primer was also found aligned within more central regions, most likely because the transcript with the primer was shorter in this species. Because of the widespread occurrence of the primer, we decided to assess its impact on the protein prediction and phylogenetic inference. Here we first detail how we removed the primer sequences from the transcriptome assemblies and the predicted proteins, and then present a robustness analysis of the phylogenetic inference.

We used Cutadapt (version 3.0) to remove primer sequences from the 5' and 3' ends of both the assembled Trinity transcripts and the predicted coding sequences (CDS) we derived from them using TransDecoder. In these trimming steps, we used the longer (28nt) version of the primer. We allowed for multiple trimming of a contig, required a minimal overlap of seven bases and allowed for 10% mismatch. Using this approach, we trimmed most of the primer, as indicated by a drastic reduction in BLAST hits. On average, 32.5% of transcripts and 14.7% of CDS were trimmed (Fig. A2 A & B). However, in some cases primer sequences could not be trimmed because they were assembled into central regions of contigs, and we decided to remove these contigs in their entirety (on average 0.9% of transcripts and 0.3% of CDS were removed, Fig. A2 C & D). Most transcripts and CDS were trimmed by the length of the "short primer" sequence, but there were also numerous cases where only fragments or concatenations of the primer had to be removed (Fig. A2 E-H).

To test whether the primer influenced the TransDecoder CDS prediction, we re-ran TransDecoder on the trimmed transcript sequences. We determined the percentage of the originally predicted proteins that were recovered using the trimmed transcripts with a BLASTP search of the new proteins against the original. As a conservative estimate, we computed the percentage of original proteins that had a match with 100% identity and >90% of the sequence and where the match arose from a protein predicted from the same assembled contig. Overall, the average recovery rate was high (94.4%, range: 87.5% – 97.1%) and, therefore, protein prediction appears robust to the presence of primer.

Next, we assessed how robust our phylogenetic inference was to the removal of primer sequences, by repeating the maximum likelihood analyses, since they are the least computationally demanding. For this we translated the trimming applied to the CDS contigs to the predicted protein sequences (PEP), making sure that this did not result in a frameshift, by removing every amino acid that contained at least one trimmed base. We then repeated the final step of the phylogenomic pipeline, starting from the final orthogroups. Specifically, we realigned orthogroups using MAFFT, trimmed them with ZORRO and concatenated them into corresponding low and high occupancy alignments (i.e. L-noPrimer and H-noPrimer). The alignments without primer were slightly longer than the original alignments and also contained a few more variable sites but were generally very similar (Table A1).

**Table A1. Characteristics of the original and protein alignments without primer.**

| Alignment metrics | L | L-noPrimer | H | H-noPrimer |
|---|---|---|---|---|
| No. genes | 8128 | 8128 | 385 | 385 |
| Alignment length | 1,687,014 | 1,711,862 | 94,625 | 95,272 |
| No. variable sites | 1,157,689 | 1,166,273 | 74,175 | 74,198 |
| No. parsimony informative sites | 934,803 | 947,840 | 63,066 | 63,461 |
| Missing data (%) | 59.3 | 59.6 | 22.9 | 23.3 |

We then repeated the L-IQ-TREE, H-IQ-TREE and C-IQ-TREE analysis with the new alignments, as described in the main text, again partitioning the concatenated alignments for each ortholog, and assigning each partition the initially inferred substitution model. We assessed topological differences between the resulting phylogenies visually and by calculating the Robinson-Foulds distance (Robinson and Foulds 1981). Further, we assessed differences in branch length as the difference in total branch length and calculated the branch site score difference following (Kuhner and Felsenstein 1994), as implemented in the function *KF.dist* in the R package phangorn (Schliep 2011).

Repeats of the L-IQ-TREE and H-IQ-TREE analyses resulted in almost identical topologies and branch lengths. In the L-IQ-TREE comparison (Fig. A3), *M.* sp. 39 was placed differently within the hypodermic clade. Interestingly, the placement of *M.* sp. 39 was one of the differences between original L-IQ-TREE and all other phylogenies (Fig. 1 and Fig. S1), and this conflict is resolved in L-noPrimer-IQ-TREE. Similarly, the H-IQ-TREE comparison (Fig. A4) also resolved some conflict, since *M. lignano* is monophyletic in H-noPrimer-IQ-TREE, in agreement with all the other phylogenies except for H-ExaBayes (Fig. S1). These changes suggest that, although removal of the primer sequence has little impact on our phylogenetic hypotheses, it does potentially reduce errors. The impact of these differences on our other analysis are negligible since L-IQ-TREE was not used for any comparative analyses and H-IQ-TREE and H-noPrimer-IQ-TREE are identical in topology once duplicate species are removed.

Compared to the L-IQ-TREE and H-IQ-TREE topology, there were a few more differences between the C-IQ-TREE and C-noPrimer-IQ-TREE phylogenies (Fig. A5). All differences were driven by uncertainties in the placement of species that were solely added to the phylogeny based on the *28S rRNA* fragment and they primarily involved short branches. This is expected since, as already mentioned, this fragment contains relatively little phylogenetic information between close relatives. Since the backbone of the C-IQ-TREE phylogenies is based on the H alignment, we would not necessarily expect any changes in topology, because the *28S rRNA* alignment remained unchanged. Differences between C-IQ-TREE and C-noPrimer-IQ-TREE thus highlight that some caution is needed when interpreting the placement of the species indicated in Fig. A5.

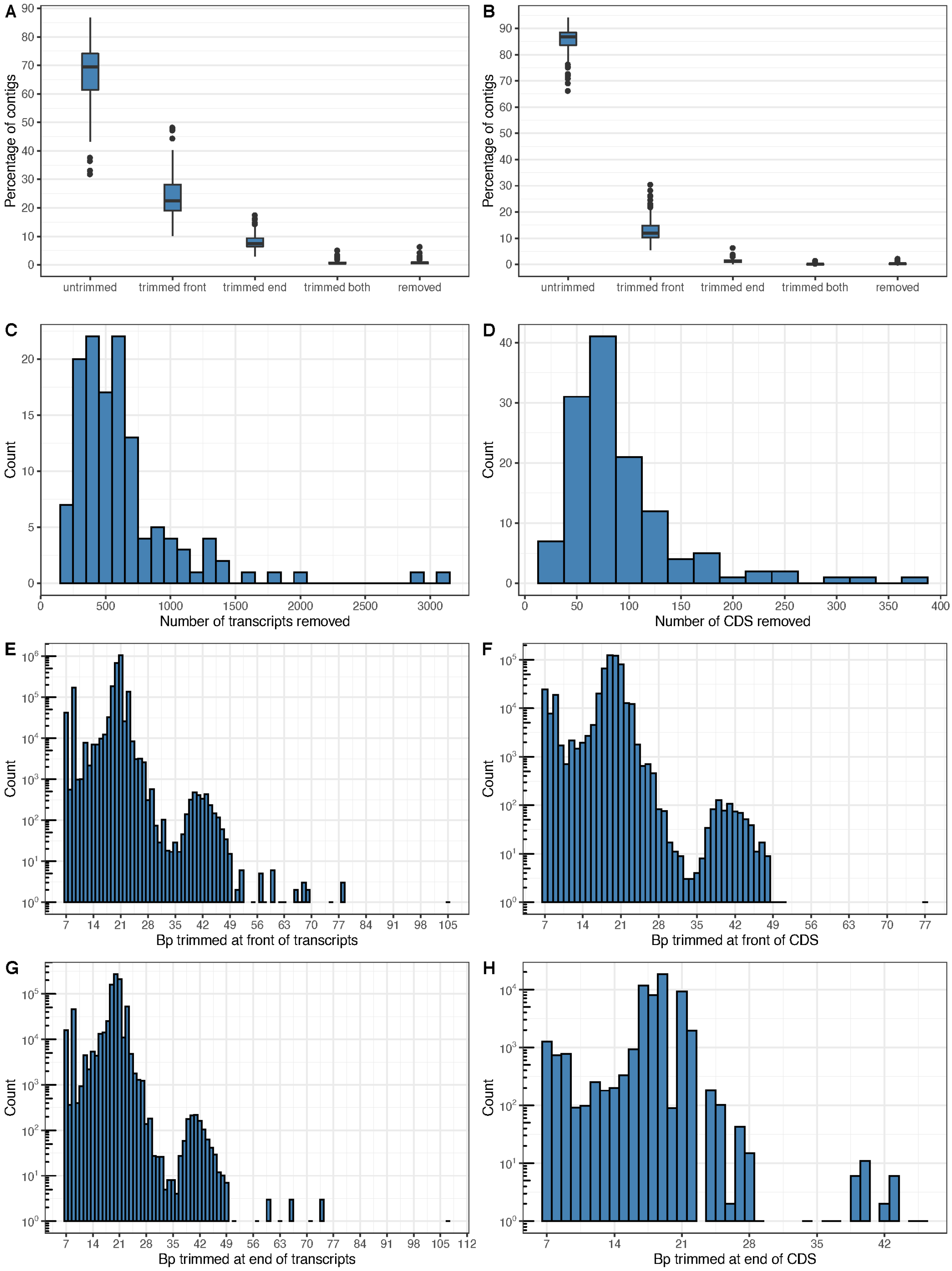


**Fig. A2. Details on primer trimming from, or the removal of, transcripts and CDS from transcriptome assemblies based RNA-Seq libraries constructed using the SMART-Seq v4 cDNA kit.** Panels on the left and right concern the raw transcriptome assemblies and TransDecoder derived predicted protein coding sequences (CDS), respectively. (A & B) Boxplots of the percentage of contigs that remained untrimmed, were trimmed (at the front, end, or both), or were removed in each transcriptome. Boxes show the second and third quartile and the whiskers extend up to 1.5 times the interquartile range. (C & D) Histograms of the number of contigs that were removed in each transcriptome due to primer in central regions. (E-H) Histograms of the number of individual transcripts across all transcriptomes that were trimmed by a certain length at the front (E & F) and/or the end (G & H). Note that the y-axis of E-H is on a log10 scale.

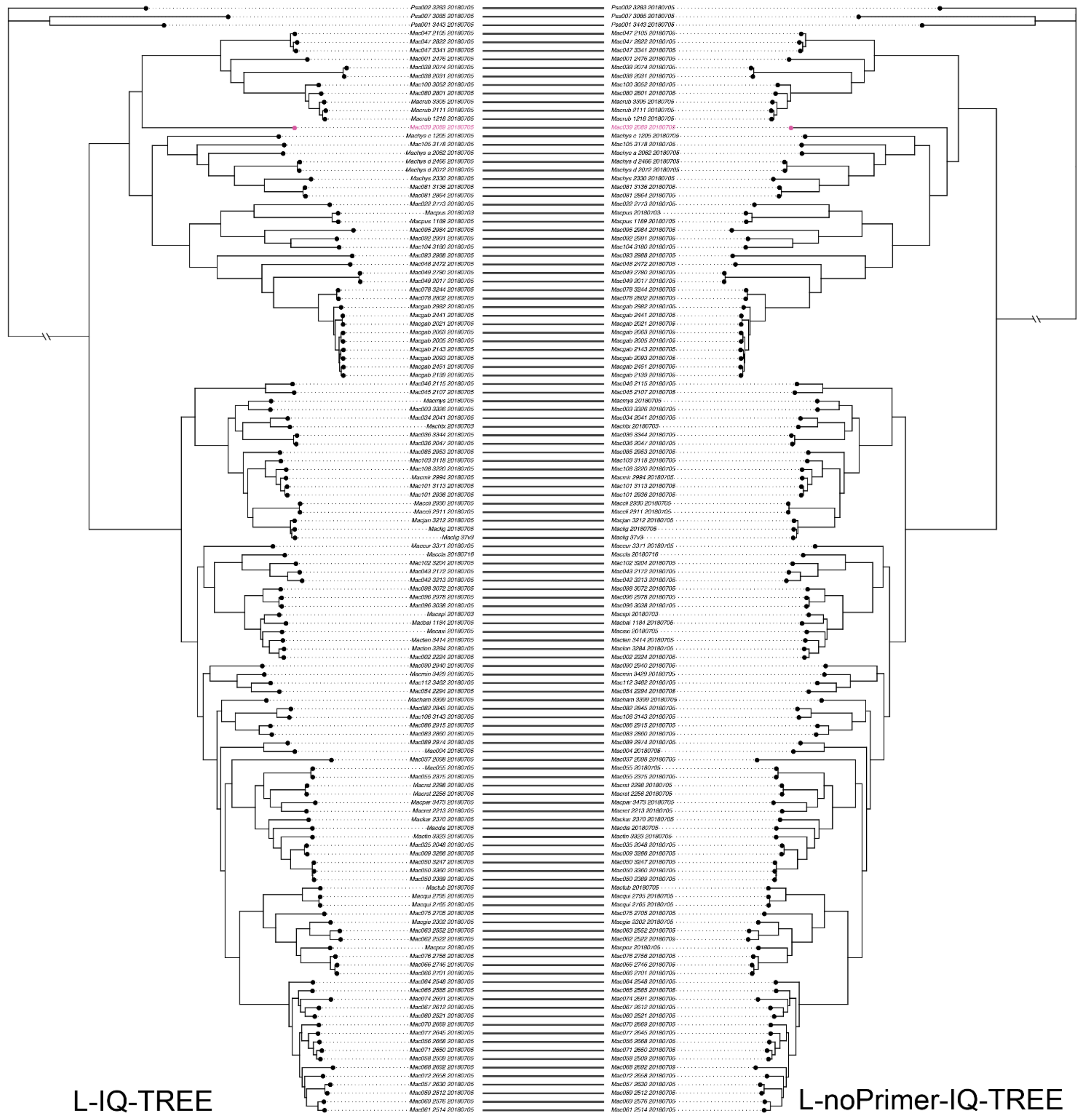


**Fig. A3. Comparison of IQ-TREE phylogenies based on the L and the L-noPrimer alignment.** The topology is identical except for the placement of *M.* sp. 39 (indicated in pink), which is more closely related to the clade containing *M. rubrocinctum* in L-IQ-TREE as opposed to being more closely related to the clade containing *M. hystricinum* in L-noPrimer-IQ-TREE (Robinson-Foulds distance: 2). Branch-length between the two trees was similar with a branch score distance of 0.043 and a 0.3% reduction in total branch length in L-noPrimer-IQ-TREE compared to L-IQ-TREE.

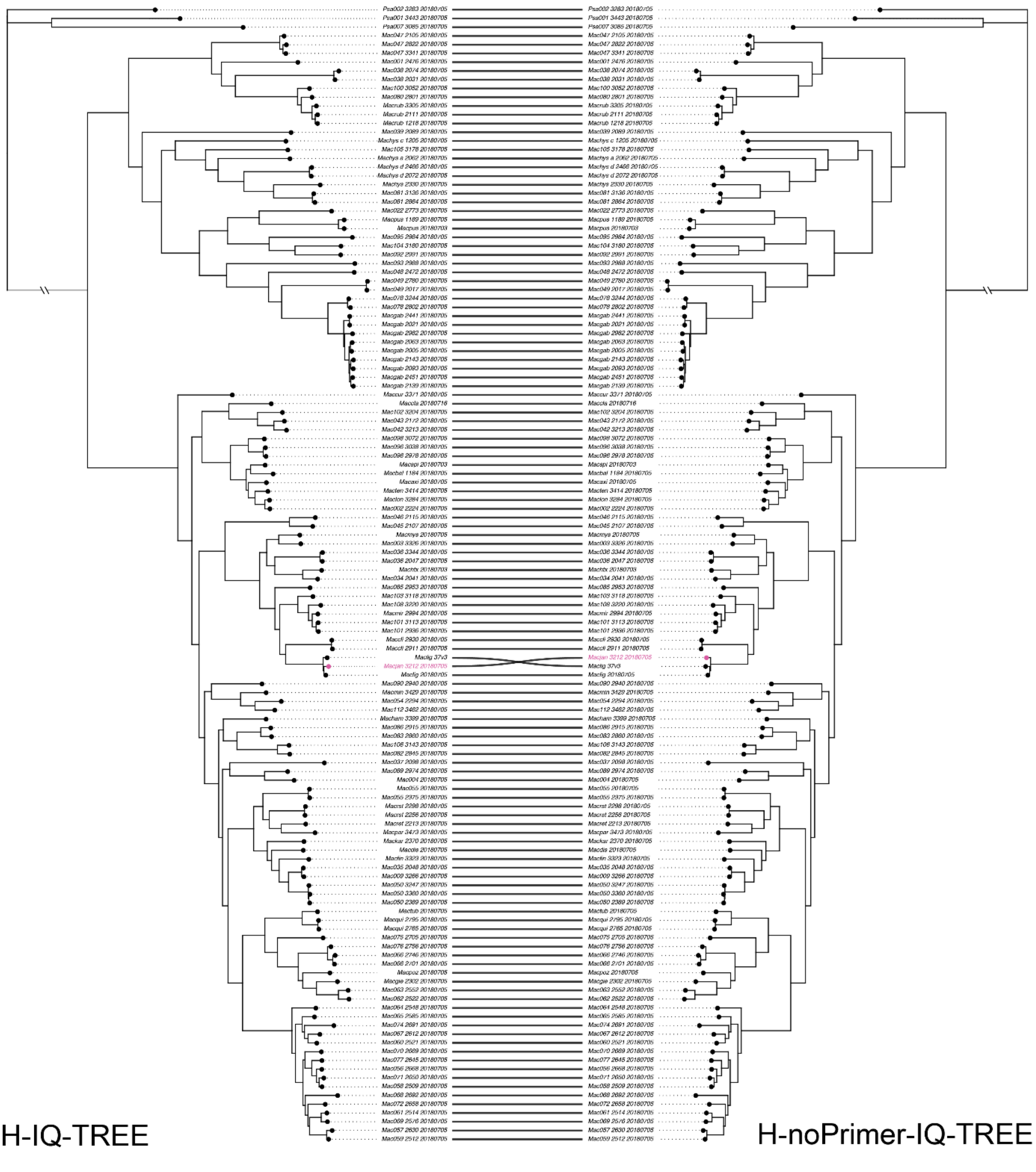


**Fig. A4. Comparison of IQ-TREE phylogenies based on the H and the H-noPrimer alignment.** The topology is identical except for the placement of *M. janickei* (indicated in pink), rendering *M. lignano* polyphyletic in H-IQ-TREE, but monophyletic in H-noPrimer-IQ-TREE (Robinson-Foulds distance: 2). Branch-length between the two trees was also similar with a branch score distance of 0.019 and a 0.9% reduction in total branch length in H-noPrimer-IQ-TREE compared to H-IQ-TREE.

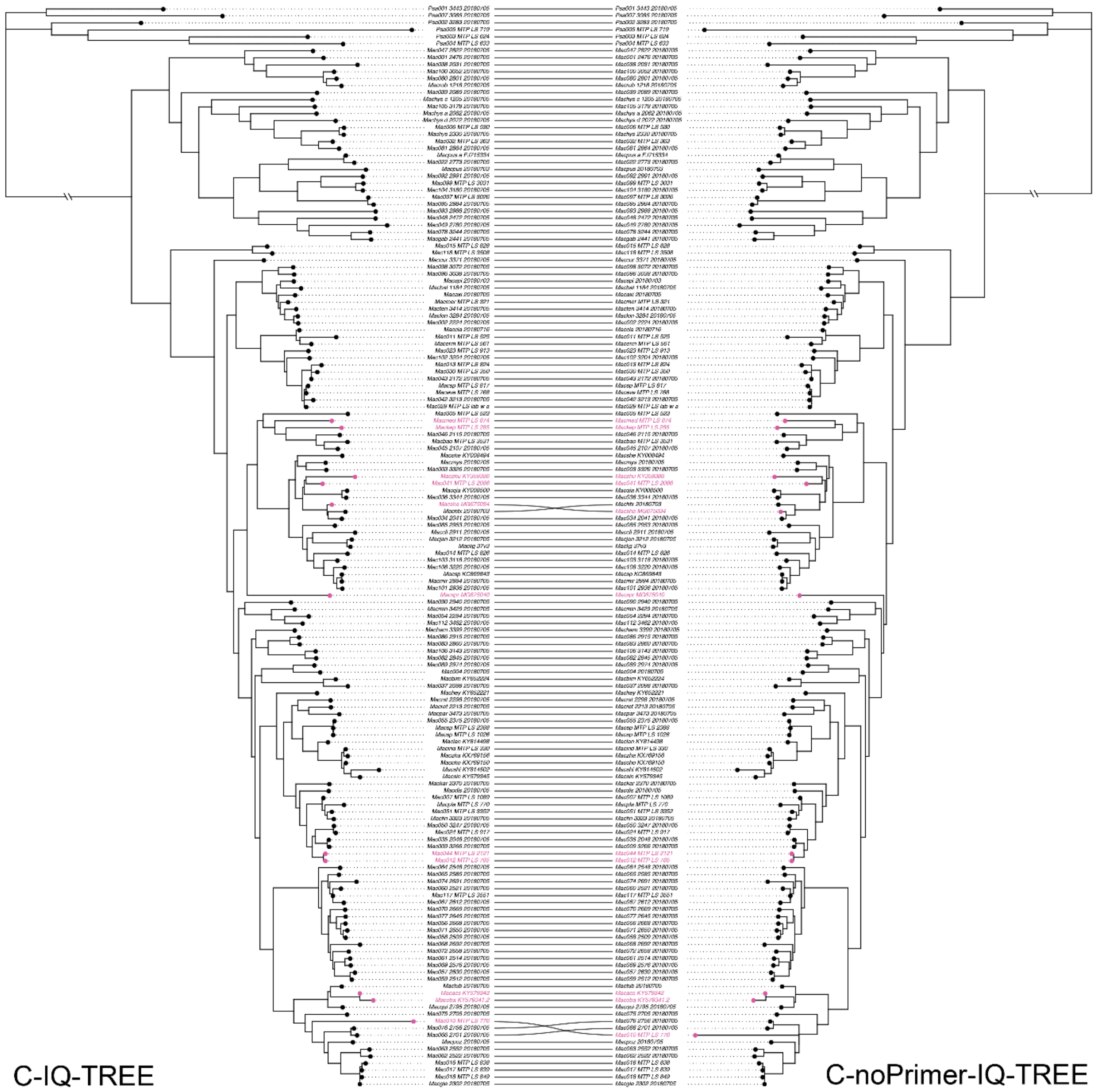


**Fig. A5. Comparison of IQ-TREE phylogenies based on the C and the C-noPrimer alignment.** The phylogeny was trimmed to remove duplicates of species for which we had more than one transcriptome since possible discrepancies based on their placement have already been assessed in Fig. A4. All differences were driven by uncertainties of the placement of species (highlighted in pink) that were solely added based on the *28S rRNA* fragment (Robinson-Foulds distance: 18). Despite these discrepancies, branch-length were similar with a branch score distance of 0.093 and a 1.32% reduction in total branch length in C-noPrimer-IQ-TREE compared to C-IQ-TREE.

### SI Species delimitation

To facilitate species delimitation we constructed haplotype networks for all 668 partial *28S rRNA* sequences included in this study (Table S1). We inferred haplotype networks for 16 groups (indicated by numbers in Fig. A6) and visualised them including the morphology of the stylet, and when diagnostic also the sperm, for each cluster. As described in the methods, we delimit species with a distance in the network >3 mutational steps. Since in some cases the networks become quite large, we severed all connections larger than 12 steps and display the separated clusters for each group. In each display (Fig. A7-A22), differences between the sequences are represented by edges in the graph, and the scale bar represents 20 µm.

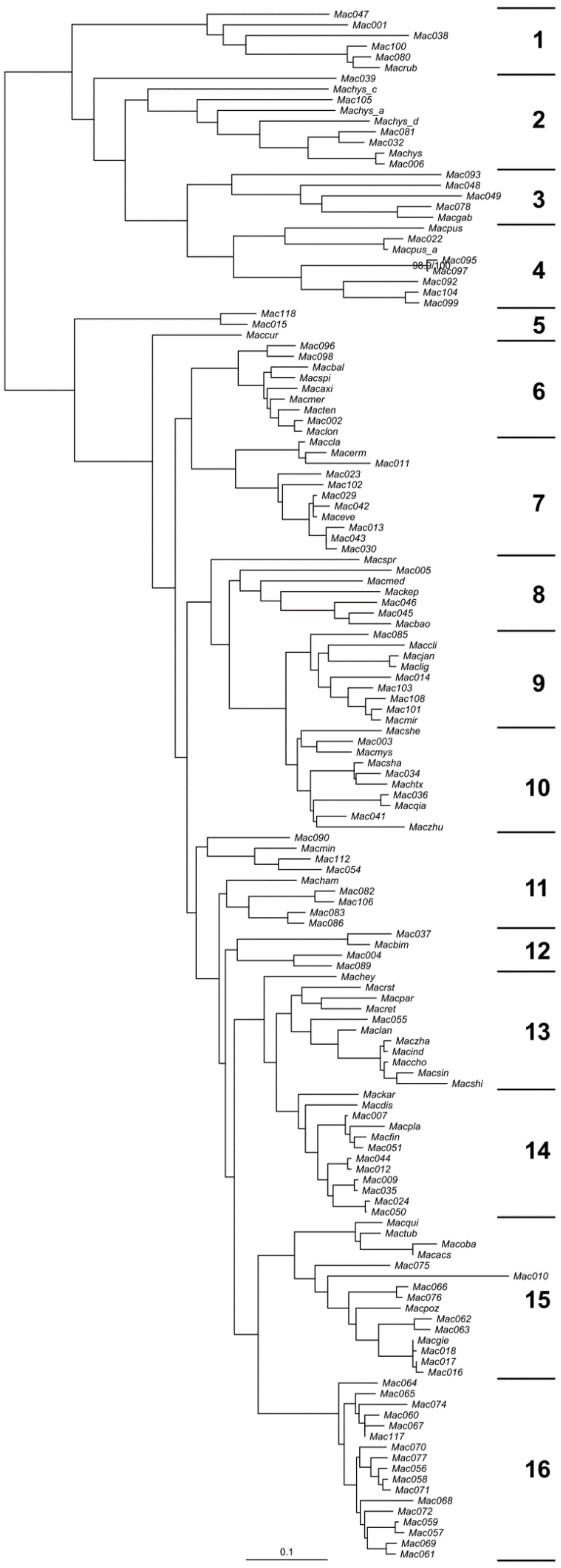


**Fig. A6. C-IQ-TREE phylogeny indicating the 16 groups for which haplotype networks were constructed.** Scale bar represents substitutions per site and large numbers denote the group number (see Fig. A7-A22).

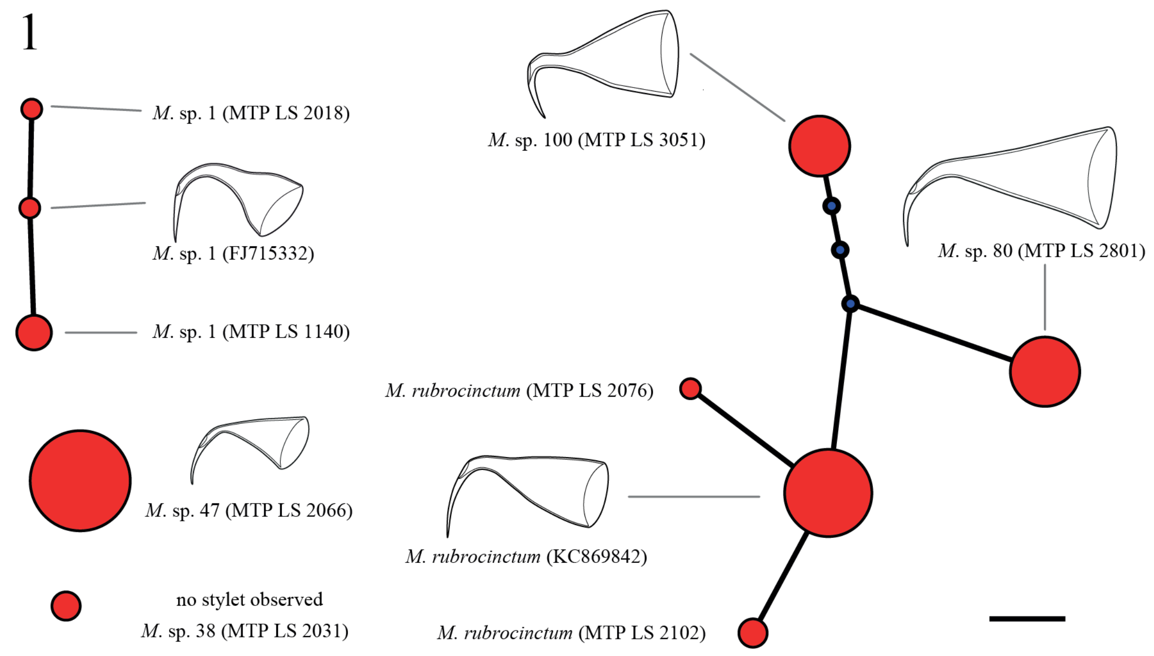


**Fig. A7. TCS haplotype network of group 1.** The size of the nodes in all haplotype networks is proportional to the number of specimens with this haplotype, and the nodes are labelled with one representative specimen. Edges represent one mutational difference and blue nodes intermediate haplotypes that were not observed. While molecularly distinct, we did not observe a stylet for *M.* sp. 38.

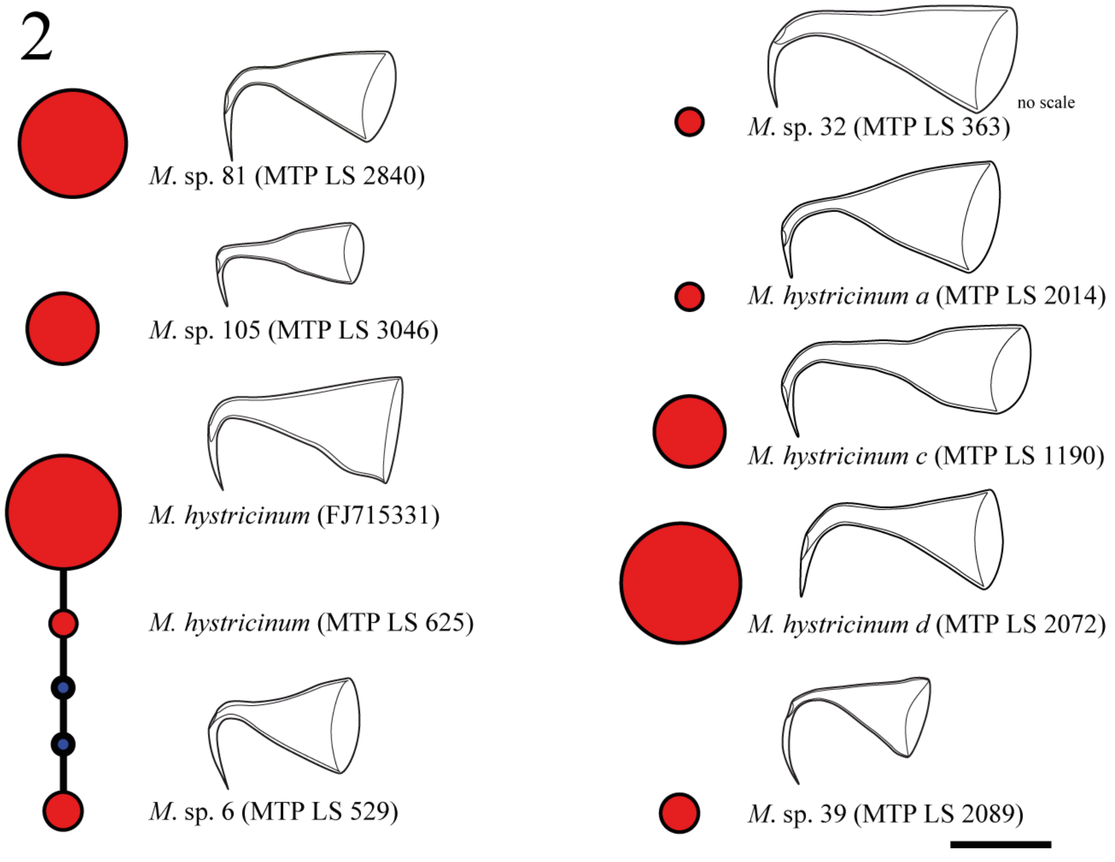


**Fig. A8. TCS haplotype network of group 2.** Note the generally high levels of divergence between these morphologically very similar species (see also Fig. A6).

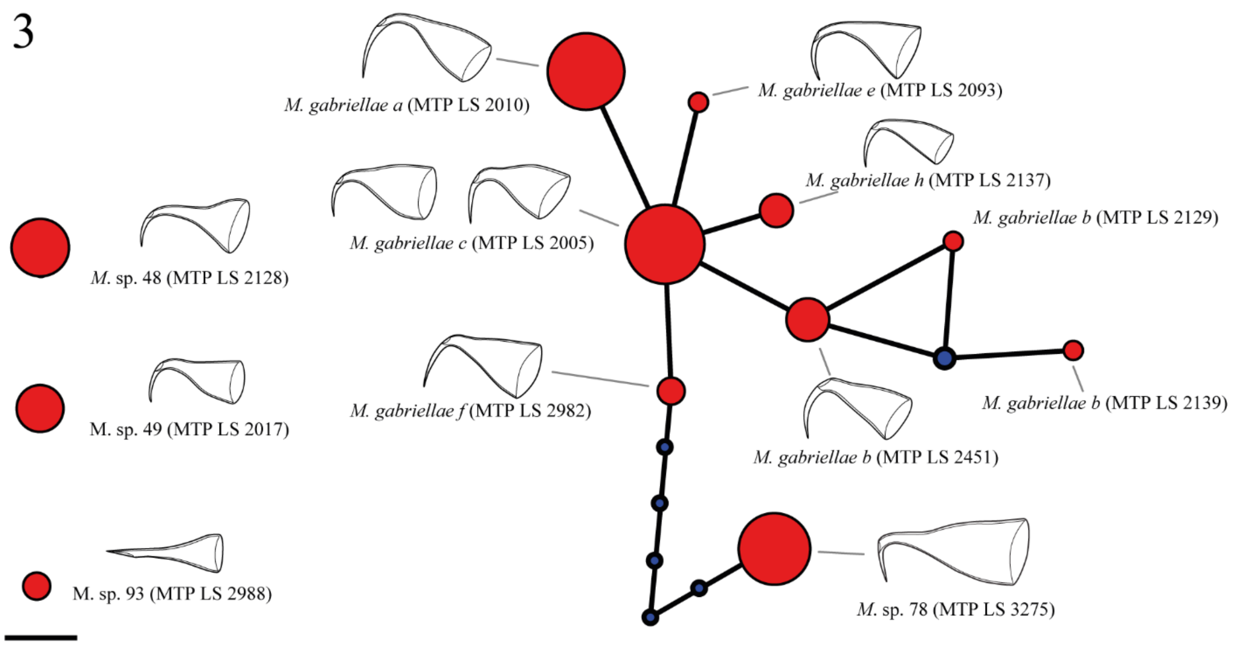


**Fig. A9. TCS haplotype network of group 3.** All haplotypes denoted with *M. gabriellae* have been grouped into one species. While some differences in the stylets are apparent in these drawings, the variation within each cluster is substantial and overlapping with the other morphologies, and hence, we have not been able to consistently assign clusters based on morphology.

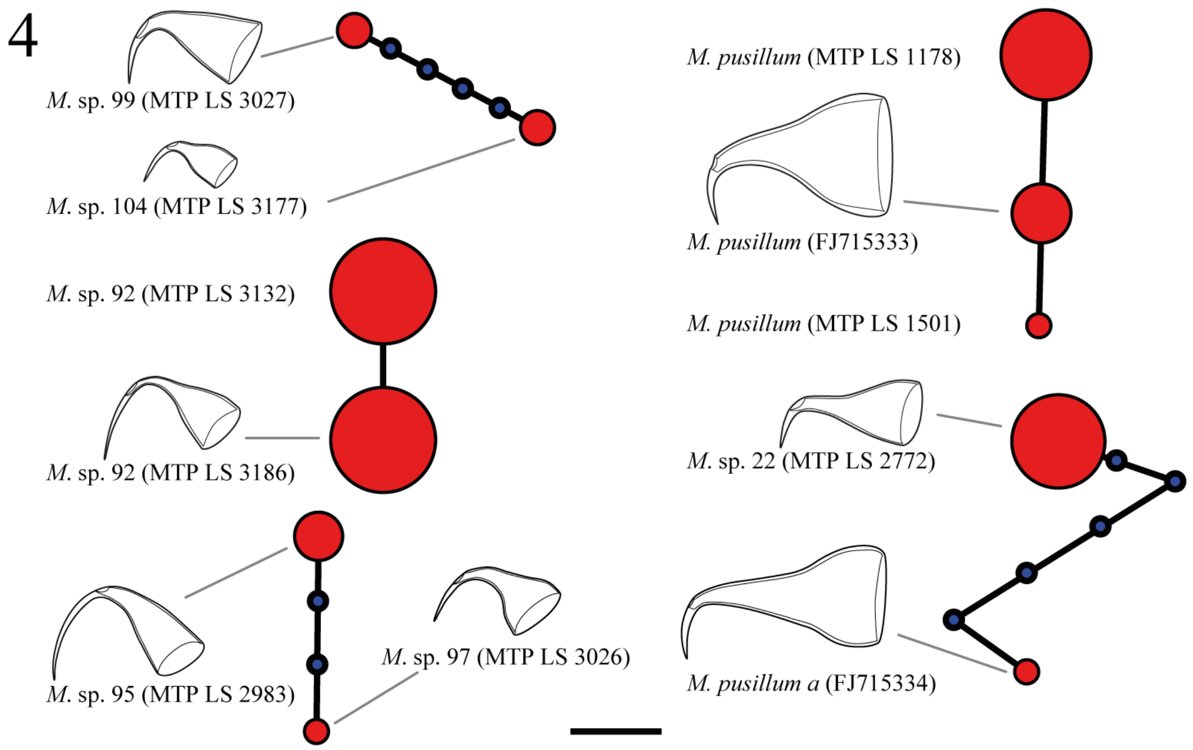


**Fig. A10. TCS haplotype network of group 4.** Note that *M. pusillum a*, which is so named because it has a close resemblance with *M. pusillum*, clusters more closely with *M.* sp. 22.

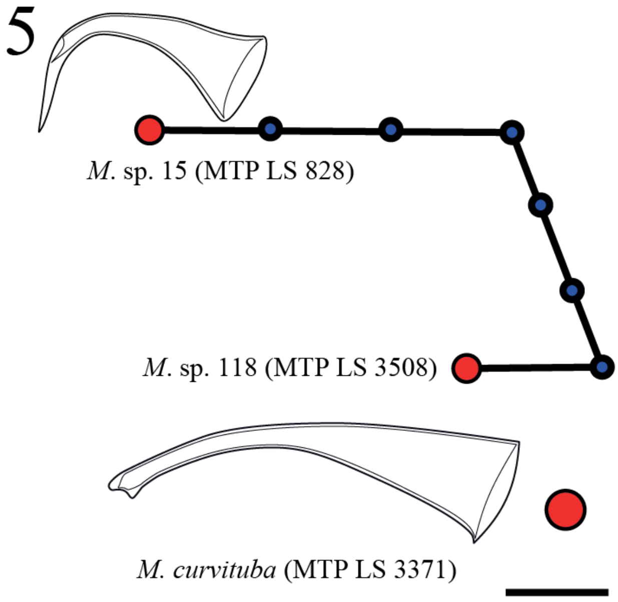


**Fig. A11. TCS haplotype network of group 5.** *M.* sp. 118 was not sexually mature when sampled, and we have no morphological information on the stylet. Also, we do not have microscope scale information for *M.* sp. 15.

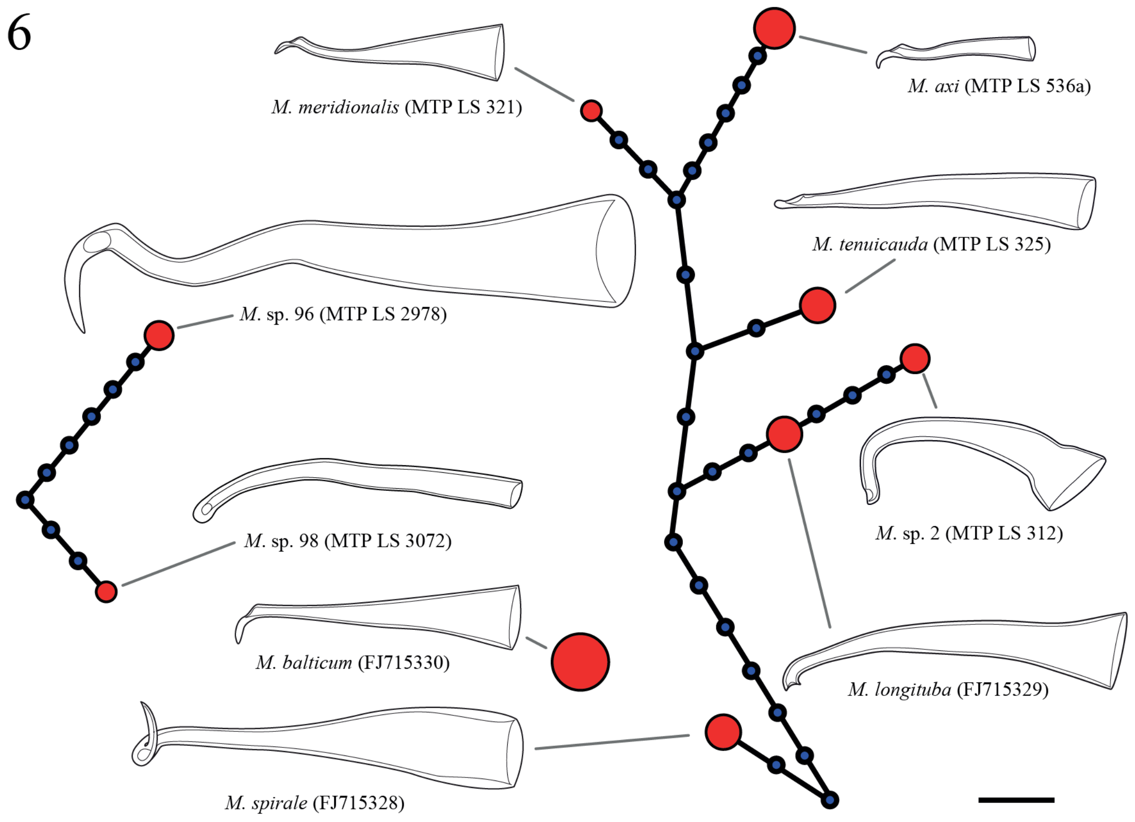


**Fig. A12. TCS haplotype network of group 6.**

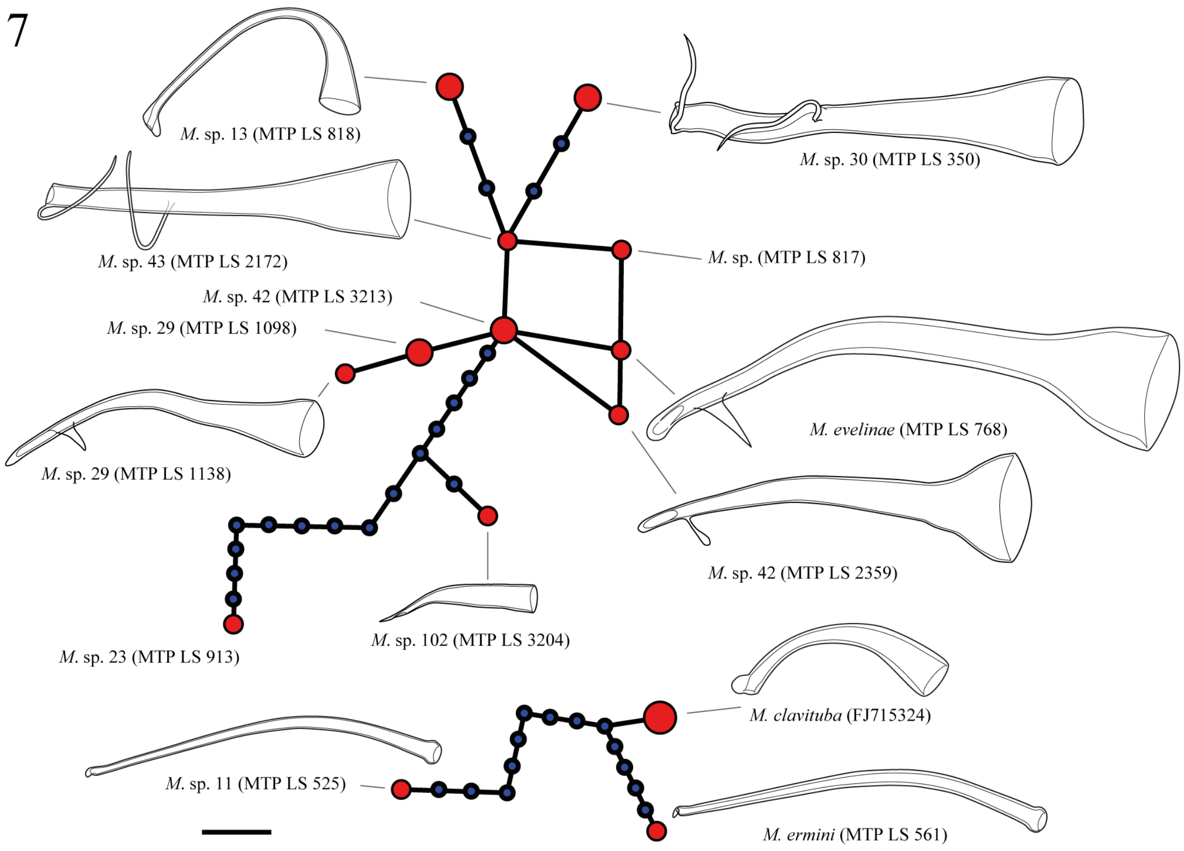


**Fig. A13. TCS haplotype network of group 7.** *M.* sp. (MTP LS 817) was not sexually mature when sampled, and we have no morphological information on the stylet. Since it clusters closely with some of the other species, we have not assigned it a provisional species number. *M.* sp. 29, *M.* sp. 42, *M.* sp. 43 and *M. evelinae* are all connected with <3 differences but are clearly distinct morphologically. Specifically, *M.* sp. 42 is distinct due to its funnel-shaped proximal opening and a rounded lateral protrusion just proximal from the opening. While *M. evelinae* is superficially similar, the stylet is larger, and the lateral protrusion is sharp. *M.* sp. 29 also has a sharp protrusion, but it is located further proximal, the distal opening is larger, and the stylet has an additional curve. *M.* sp. 43 bears little resemblance to the other three, having a straight stylet that has two thin lateral protrusions.

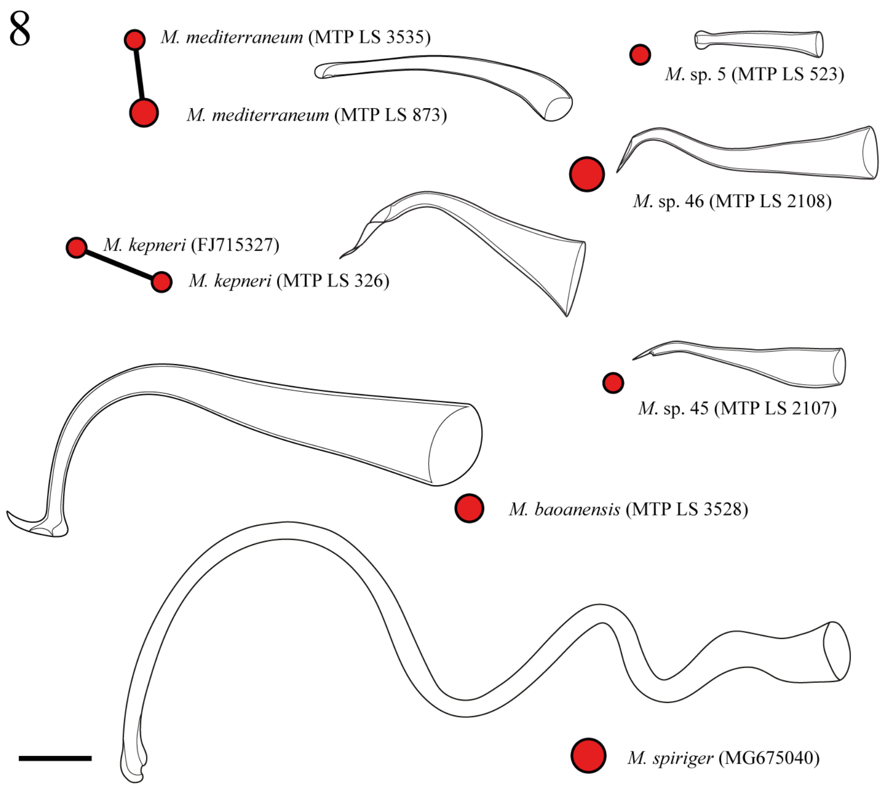


**Figure A14. TCS haplotype network of group 8.**

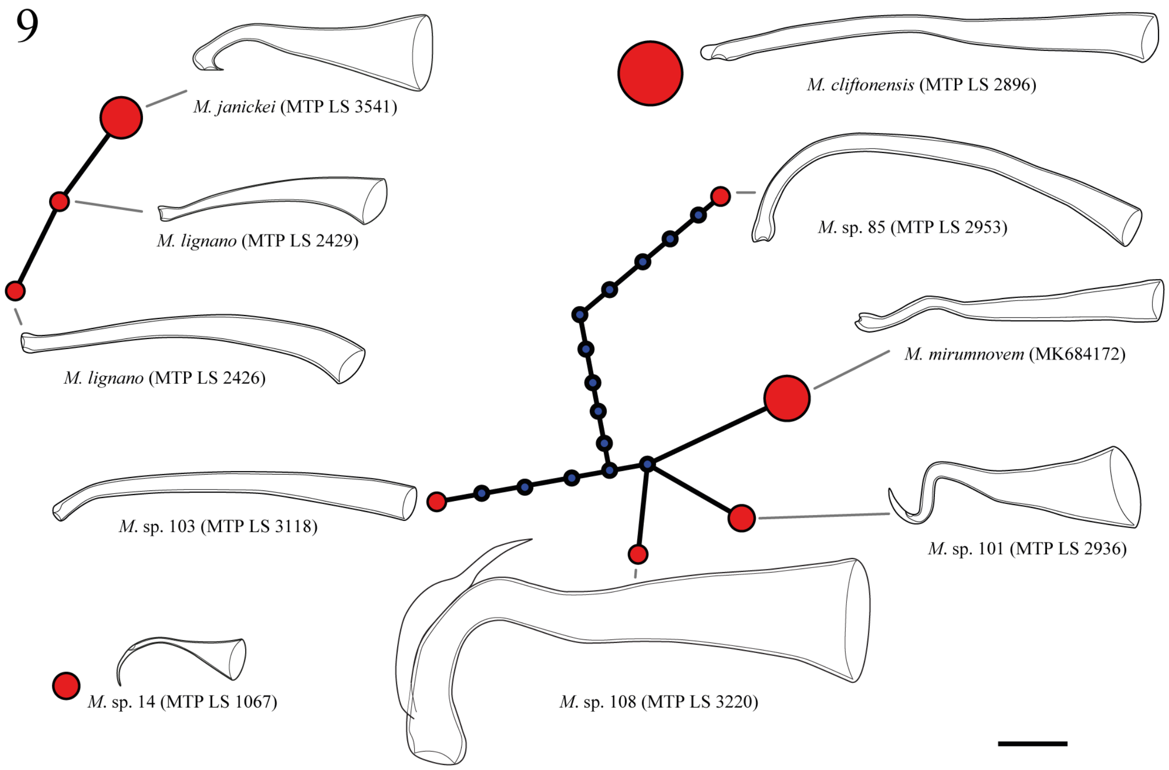


**Fig. A15. TCS haplotype network of group 9.** The specimen MTP LS 2429 represents the *M. lignano* population from Italy, while MTP LS 2426 represents a population from Greece, which has a larger stylet. However, the shape variation within the former can include morphologies like the one drawn for the latter. Note that our *28S rRNA* sequence of *M. mirumnovem* is identical to GenBank accession KC869843 of a *Macrostomum* sp. from South Australia (Laumer and Giribet 2014). *M. mirumnovem*, *M.* sp. 101, and *M.* sp. 108 are clearly distinct, with much larger size in *M.* sp. 108 and a sharp twisted distal region in *M.* sp. 101.

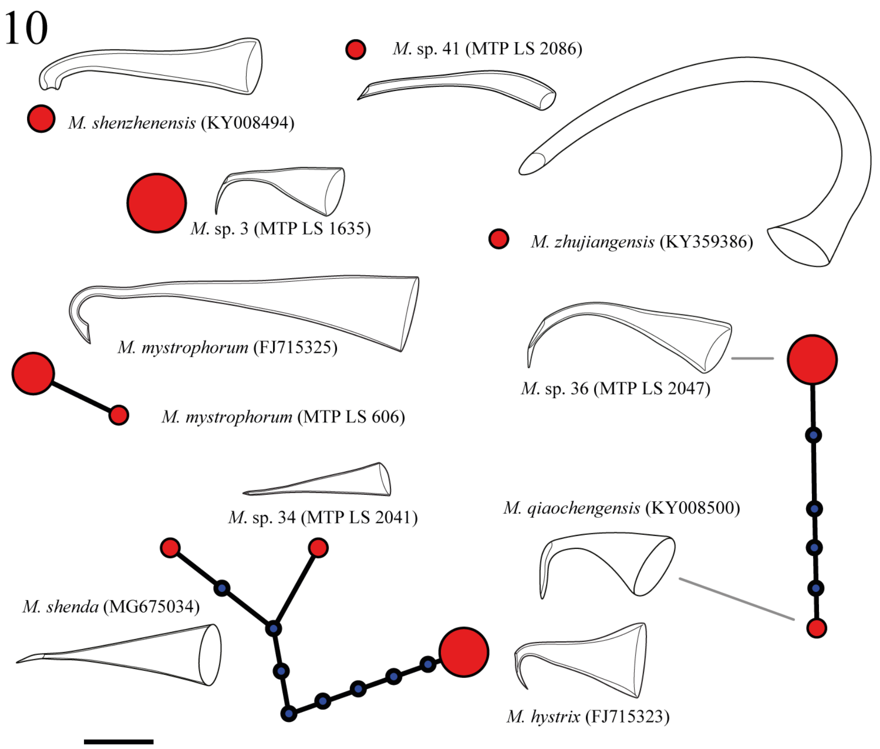


**Fig. A16. TCS haplotype network of group 10.** *M.* sp. 34 and *M. shenda* are similar in stylet morphology and general habitus, however, in *M. shenda* the stylet is larger.

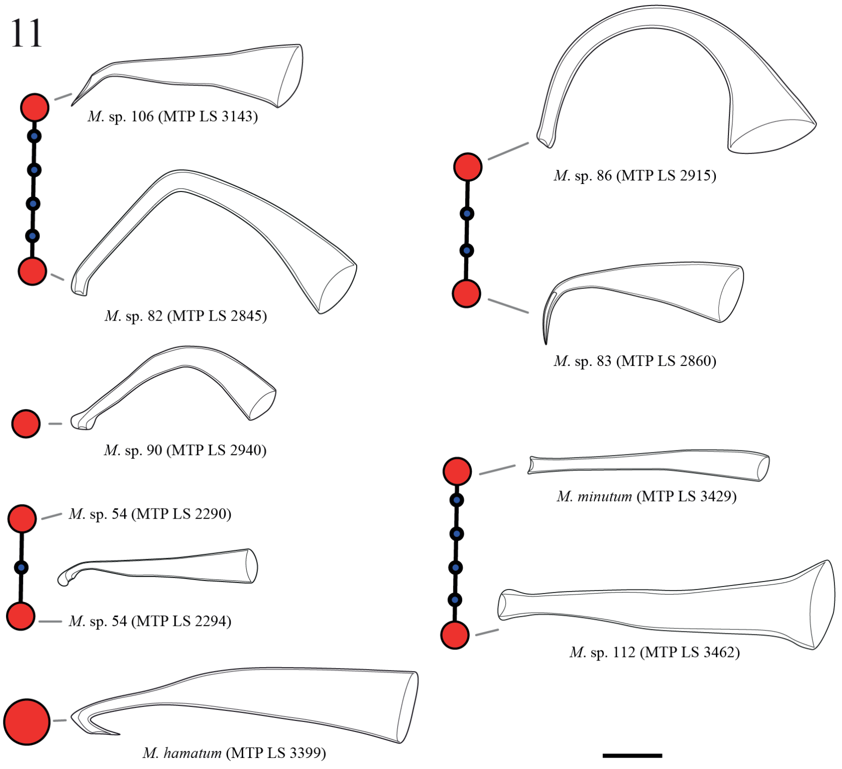


**Fig. A17. TCS haplotype network of group 11**. The stylet of *M. minutum* and *M.* sp. 112 differ in that the former is longer, wider, and has a clear distal blunt thickening, while the latter does not have this thickening and does not have a funnel-shaped proximal opening.

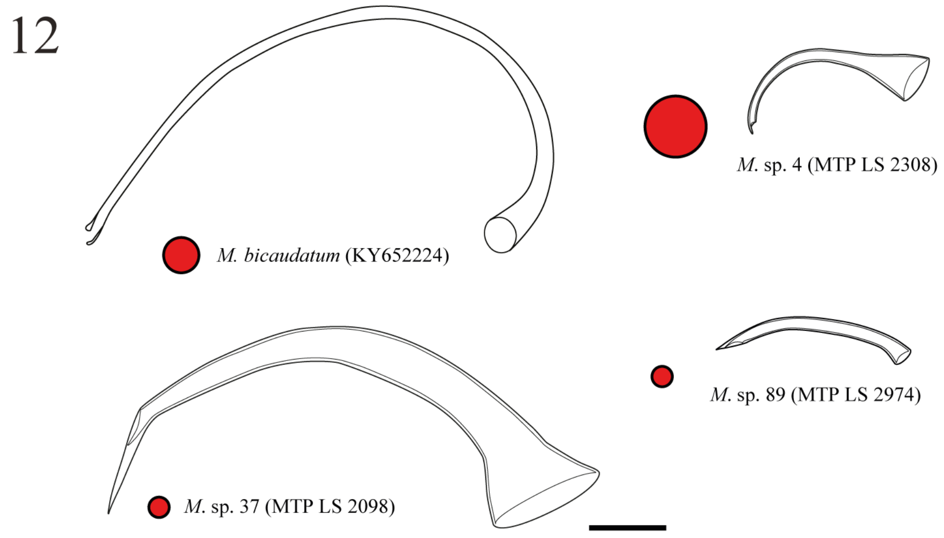


**Fig. A18. TCS haplotype network of group 12.**

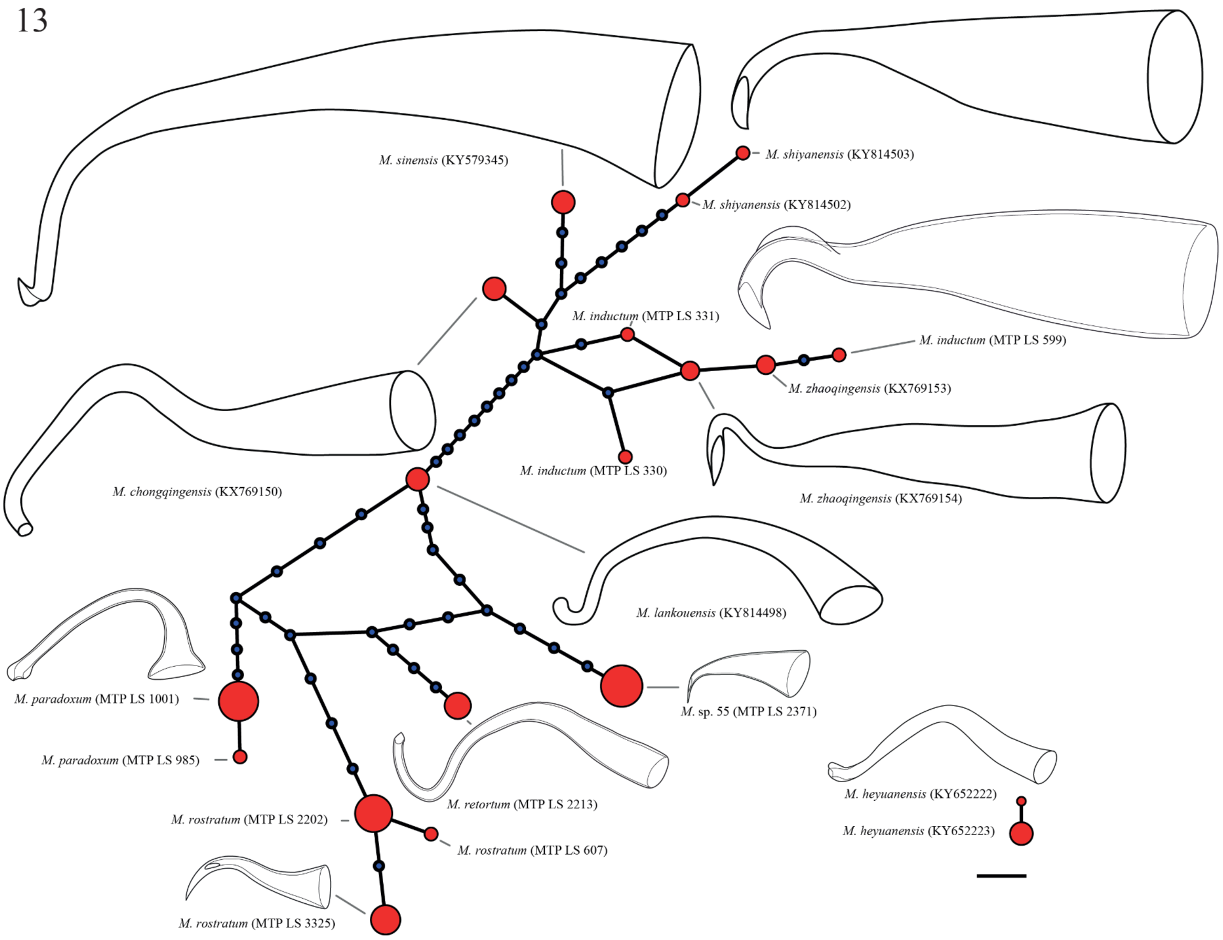


**Fig. A19. TCS haplotype network of group 13.** Our *M. inductum* samples cluster closely with *M. zhaoqingensis*, and they are also very similar in stylet and sperm morphology. The latter could thus be a re-description of the former and should be further investigated taxonomically.

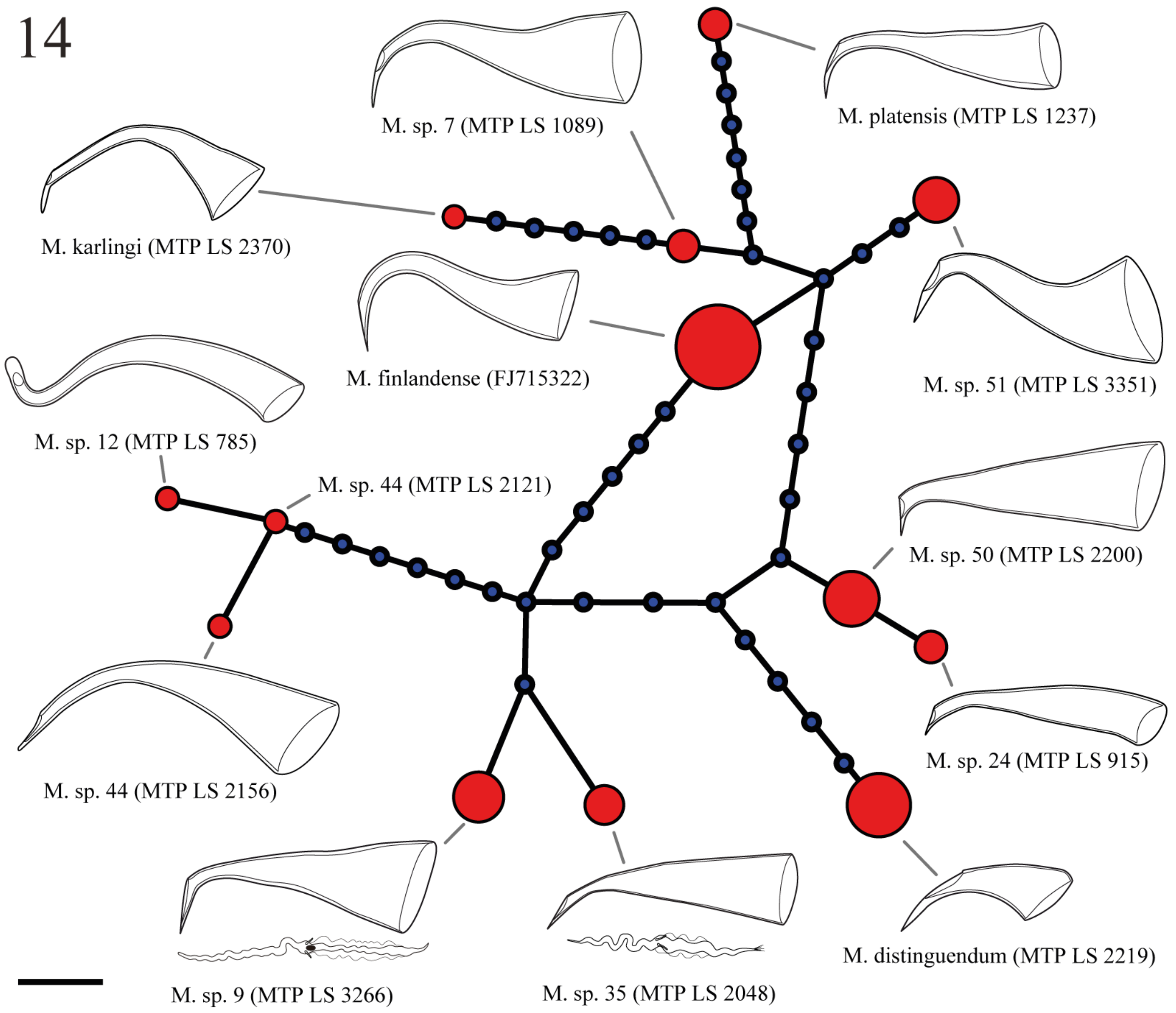


**Fig. A20. TCS haplotype network of group 14.** Stylets are similar in shape between *M.* sp. 9 and *M.* sp. 35, but with a long drawn out distal tip and a broader base in the former. The sperm between the two species is diagnostic, with a clearly visible dense body close to the bristles in *M.* sp. 9. This structure, which could be the nucleus, is not visible in *M.* sp. 35. Moreover, the sperm of *M.* sp. 35 has a brush. The stylets of *M.* sp. 12 and *M.* sp. 44 differ in the width of the base, being much wider in the latter and ending in an oblique cut, while there is an additional curve in the former ending in a blunted subterminal opening. The stylets of *M.* sp. 24 and *M.* sp. 50 differ in the width of the base, being much wider in the latter, with a more pronounced 90° turn of the distal thickening.

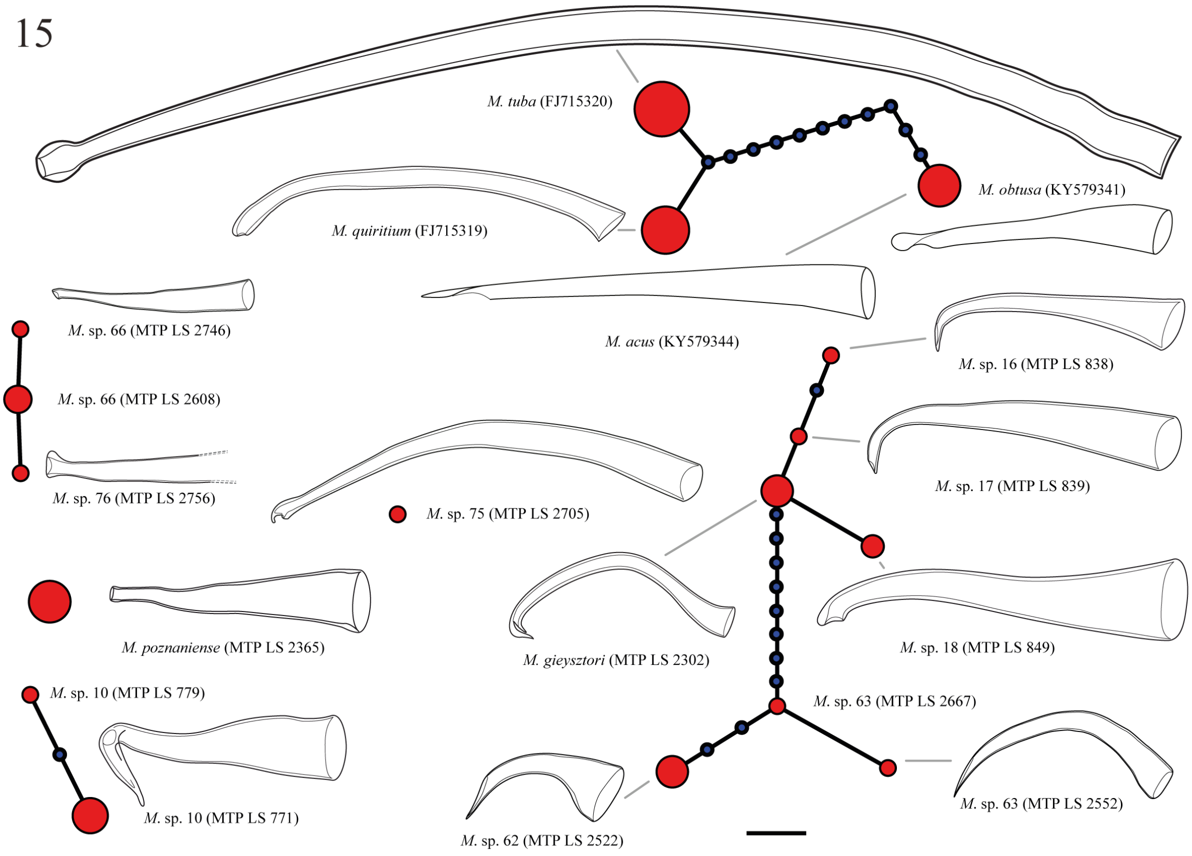


**Fig. A21. TCS haplotype network of group 15.** The stylet of *M*. sp. 16 is shorter than *M*. sp. 17 and the distal curve of the stylet is sharper in the former ending in a strongly oblique tip, while the curve is gentler in the latter and ends much less oblique. *M*. sp. 18 differs from the former two since it has a longer wider stylet that terminates subterminally and has a slight thickening on the convex part of the distal curve. All three species differ from *M. gieysztori*, which has a thinner stylet with a pronounced additional curve and a subterminal opening pointing proximally and a clear anterior thickening.

16

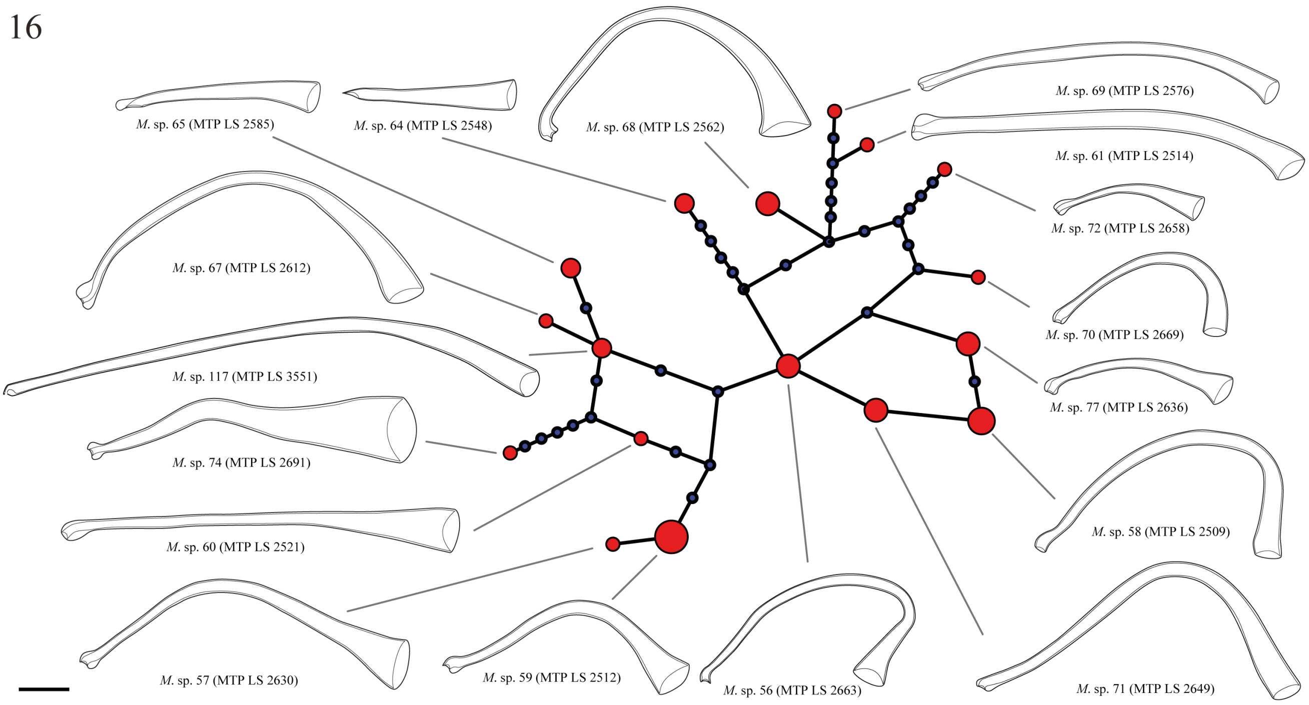


**Fig. A22. TCS haplotype network of group 16.** All species were collected in Lake Tanganyika, except for *M*. sp. 117, which was collected from Lake Malawi. Some species cluster closely in *28S rRNA*. *M*. sp. 65, *M*. sp. 67 and *M*. sp. 117 can easily be distinguished. *M*. sp. 61 differs from *M*. sp. 69 in that the latter has a more strongly pronounced distal thickening. *M*. sp. 56 differs from *M*. sp. 58 in that the proximal opening is pointing towards the distal opening in the former. *M*. sp. 71 is distinct from both in that the curve of the stylet is much more open, and the distal opening has a slight thickening on the concave side.
