## Supplementary figures and images for "Large-scale phylogenomics of the genus *Macrostomum* (Platyhelminthes) reveals cryptic diversity and novel sexual traits"

### Figure S1

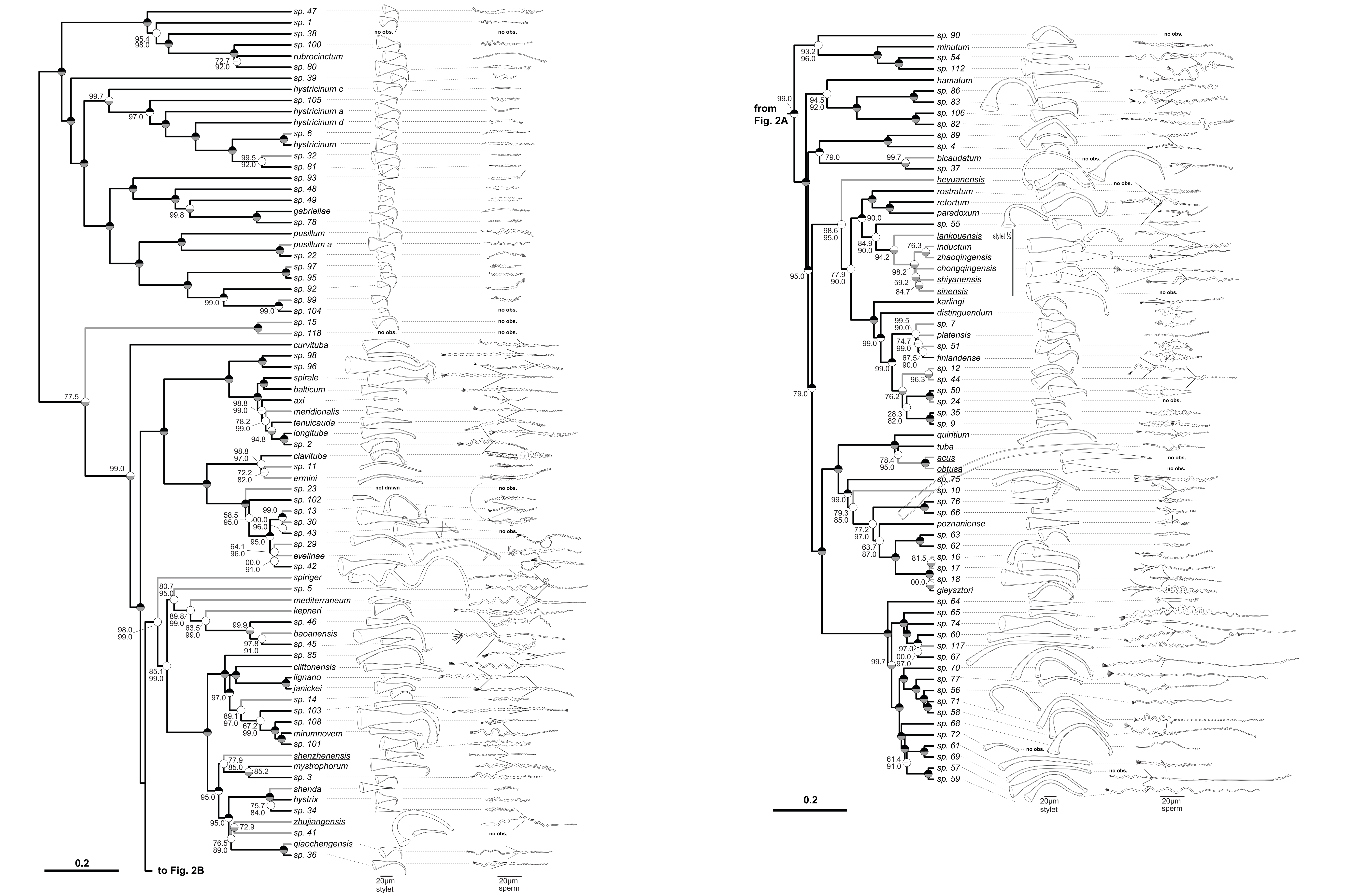

### Figure S2

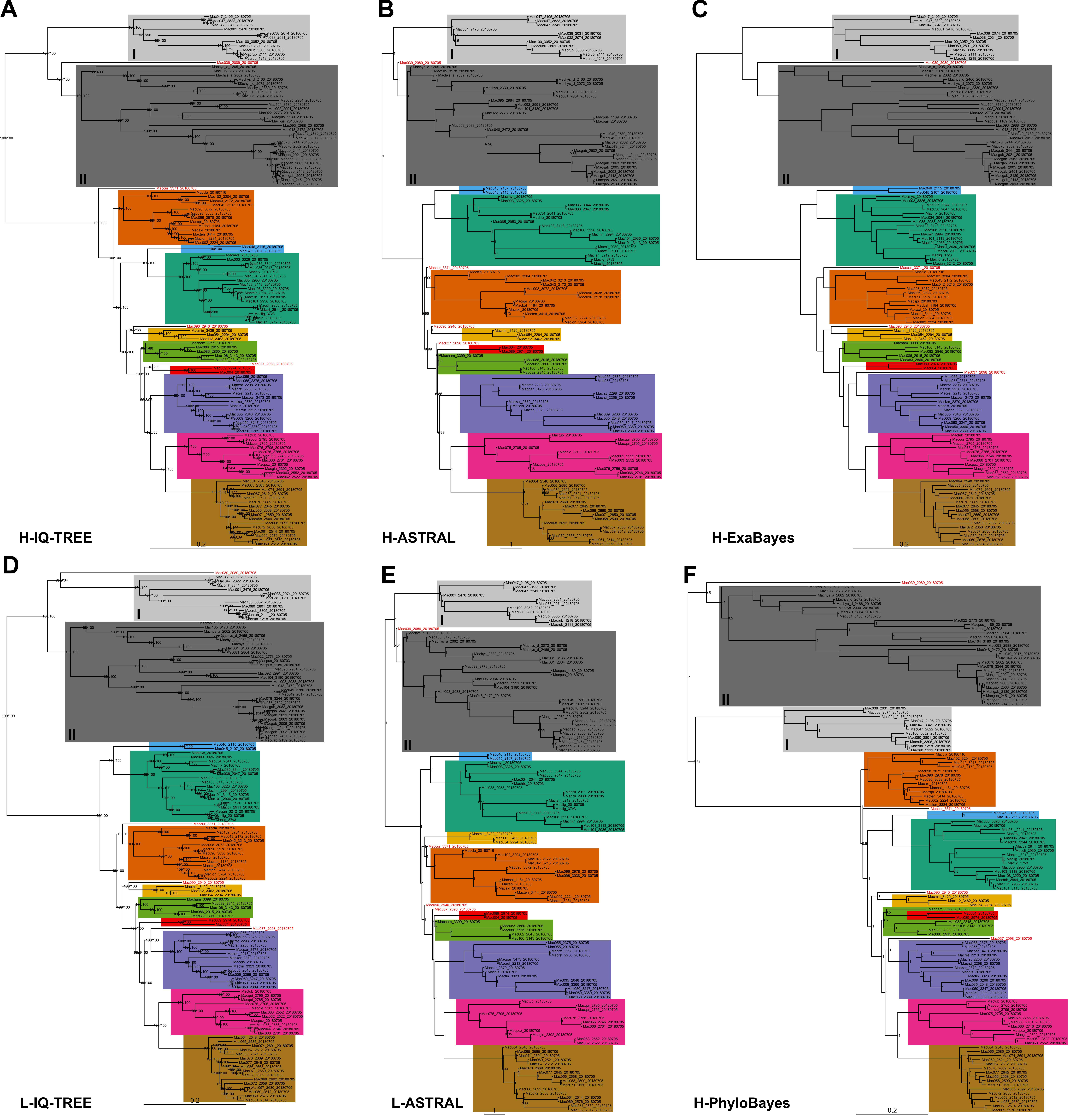

### Figure S3

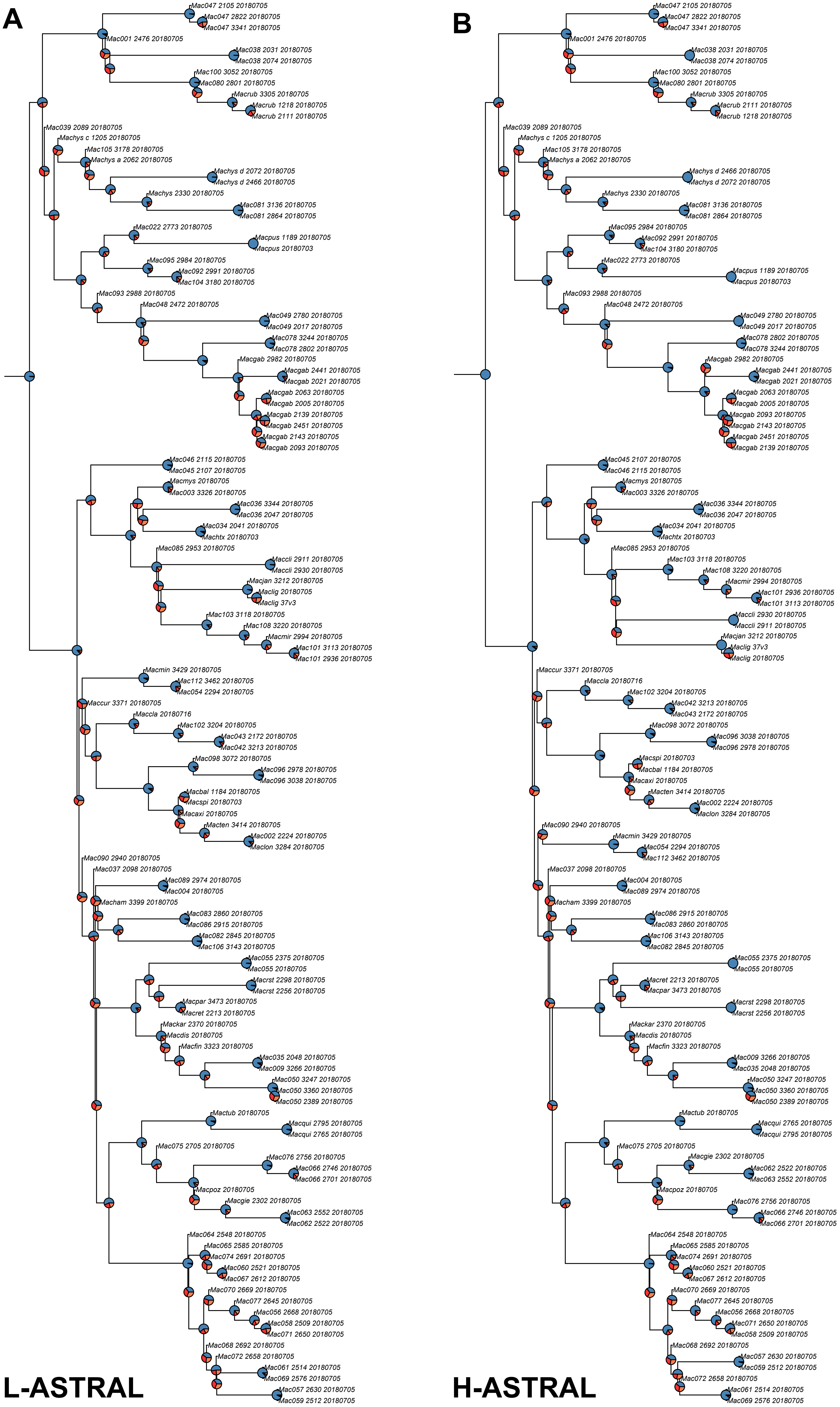
